# Diversity of photosynthetic picoeukaryotes in eutrophic shallow lakes as assessed by combining flow cytometry cell-sorting and high throughput sequencing

**DOI:** 10.1101/551598

**Authors:** Sebastián Metz, Adriana Lopes dos Santos, Manuel Castro Berman, Estelle Bigeard, Magdalena Licursi, Fabrice Not, Enrique Lara, Fernando Unrein

**Affiliations:** Instituto de Investigaciones Biotecnológicas-Instituto Tecnológico de Chascomús (IIB-INTECH), UNSAM-CONICET. Av. Intendente Marino Km 8,200, (7130) Chascomús, Buenos Aires, Argentina; Sorbonne Université, CNRS, Laboratoire Adaptation et Diversité en Milieu Marin UMR7144, Station Biologique de Roscoff, 29680 Roscoff, France; Instituto Nacional de Limnología (INALI), CONICET-UNL. Ciudad Universitaria - Paraje el Pozo s/n (3000), Santa Fé, Argentina.; Laboratory of Soil Biodiversity, University of Neuchâtel, Rue Emile Argand 11, CH-2000 Neuchâtel, Switzerland; Present address: Real Jardín Botánico de Madrid, CSIC, Plaza de Murillo 2, 28014 Madrid, Spain; Asian School of the Environment, Nanyang Technological University, 50 Nanyang Avenue, Singapore 639798

**Keywords:** picoplankton, eutrophic shallow lakes, Flow cytometry, Metabarcoding

## Abstract

Photosynthetic picoeukaryotes (PPE) are key components of primary production in marine and freshwater ecosystems. In contrast with those of marine environments, freshwater PPE groups have received little attention. In this work, we used flow cytometry cell sorting, microscopy and metabarcoding to investigate the composition of small photosynthetic eukaryote communities from six eutrophic shallow lakes in South America, Argentina. We compared the total molecular diversity obtained from PPE sorted populations as well as from filtered total plankton samples (FTP). Most reads obtained from sorted populations belonged to the classes: Trebouxiophyceae, Chlorophyceae and Bacillariophyceae. We retrieved sequences from non-photosynthetic groups, such as Chytridiomycetes and Ichthyosporea which contain a number of described parasites, indicating that these organisms were probably in association with the autotrophic cells sorted. Dominant groups among sorted PPEs were poorly represented in FTP and their richness was on average lower than in the sorted samples. A significant number of operational taxonomic units (OTUs) were exclusively found in sorting samples, emphasizing that sequences from FTP underestimate the diversity of PPE. Moreover, 22% of the OTUs found among the dominant groups had a low similarity (<95%) with reported sequences in public databases, demonstrating a high potential for novel diversity in these lakes.

## INTRODUCTION

Photosynthetic picoeukaryotes (PPEs) are key components of planktonic food webs and contribute significantly to phytoplankton biomass and primary production in aquatic systems (Li 1994; Callieri 2008). To better understand the functioning of these ecosystems, it is of key importance to document the diversity and community structure of aquatic microorganisms. Due to their small size, most PPE groups lack of conspicuous morphological characters, which precludes their precise taxonomic identification with classical light microscopy (Potter and Lajeunesse 1997). Since the early 2 000’s, molecular methods have proved their potential to reveal unpredicted diversity of PPE in marine waters (Díez, Pedrós-Alió and Massana 2001; Moon-Van Der Staay 2001; Not *et al.* 2004). These seminal works showed that Prasinophytes, Haptophytes and Pelagophyceae groups represented a significant proportion of PPE whose importance has been highlighted during the last years (Jardillier *et al.* 2010; Worden *et al.* 2012; Gómez-Pereira *et al.* 2013; Unrein *et al.* 2014; Lopes dos Santos *et al.* 2017).

As compared to their marine counterparts, the diversity of freshwater eukaryotic plankton has been far less investigated. A fundamental ecological difference with marine systems resides in the fact that lakes and rivers are physico-chemically way more heterogeneous than the oceans (Simon *et al.* 2015). Continental waters do not have a continuous geographical distribution, which favours local endemism (Schiaffino *et al.* 2016; Fernandez *et al.* 2017) and therefore may lead to a greater diversity of organisms. Debroas *et al.* (2017) predicted that freshwater microbial eukaryotic richness would be around 200 000 – 250 000 species, comparing to oceans where estimated eukaryotic ribosomal diversity saturate at ca. 150 000 operational taxonomic units (OTUs) (de Vargas *et al.* 2015). As far as these extrapolations can be trusted, the few studies that addressed specifically freshwater planktonic eukaryotic diversity showed a great diversity of PPE (Lefranc et al., 2005; Lepère *et al.* 2013; Taib *et al.* 2013; Lara *et al.* 2015; Simon *et al.* 2015; Li *et al.* 2017).

Studying PPE size class with molecular approaches is not trivial. Typically, in order to remove larger eukaryotes, water samples are filtered through a 3 µm filter and the PPE biomass is collected onto a 0.2 µm filter. Such filtered samples can be used for downstream applications such as nucleic acids extraction. Early analysis of 18S rRNA gene diversity from PPE fraction recovered many sequences assigned to heterotrophic groups (Moon-van der Staay *et al.* 2001) in particular heterotrophic stramenopiles (Massana *et al.* 2004) and alveolates (Guillou *et al.* 2008) such as Syndiniales known parasites of larger phytoplankton species (Chambouvet *et al.* 2008). Larger, non-picoplanktonic organisms are also regularly detected (López-García *et al.* 2001; Massana *et al.* 2015; Simon *et al.* 2015; Schiaffino *et al.* 2016). To overcome these difficulties, the implementation of a flow cytometry cell-sorting step prior to sequencing is a good strategy to study specifically PPE (Shi *et al.* 2009; Marie *et al.* 2010). By combining flow cytometer cell-sorting and sequencing, Shi *et al*. (2009) confirmed that marine PPE are highly diversified and found unknown groups which remain uncultured until today (Tragin *et al.* 2016).

This approach offers the possibility of obtaining a combination of qualitative and quantitative data, as well as to focus on the diversity of photosynthetic organisms or their symbionts (Thompson *et al.* 2012; Gérikas *et al.* 2018).

The Pampean region is one of the most extensive plain in the world, harbouring thousands of shallow eutrophic lakes (Quirós *et al.* 2006). PPE abundance is high in these aquatic systems, reaching up to 1,9 10^5^ cells ml^−1^ (Allende *et al.* 2009; Torremorell *et al.* 2015), however, the diversity and composition of PPEs is completely unknown.

The aim of the present study was to investigate the composition of PPE communities in six eutrophic shallow lakes located in the warm-temperate Pampean region of Argentina using a combination of microscopy, flow cytometry sorting and high throughput sequencing of the V4 region of 18S rRNA gene. In order to evaluate the efficiency of cell-sorting, we also compared our results with those obtained by sequencing whole plankton community from filtered samples.

## MATERIALS AND METHODS

### Study sites and sampling

Six eutrophic shallow lakes with comparable limnological features from Pampa region were sampled between austral spring and summer of 2015 (Suppl. Table 1). The Pampa region is an extensive plain of about 800 000 km^2^ that covers the centre East of Argentina in South America (33–39°S, 57–66°W) (Giraut *et al.* 2007). The region has a warm-temperate climate with an alternation of wet and dry periods (Sierra, Hurtado and Specha 1994). Annual mean temperature and precipitation are 15.3°C and 935 mm, respectively (Iriondo and Drago 2004). The salinity of the Pampean lakes is highly variable (Quirós 2005). Accordingly to their salinity, they can be classified from oligo to meso-haline (Ringuelet 1962). Lakes Chascomús (CH), Vitel (VI) and La Limpia (LI) are located in the Salado river basin. Los Quilmes (QU) and La Juanita (LJ) are approximately 400 km away towards the west and southwest respectively. Mar Chiquita (MA2) is an albufera (lagoon) connected to the Atlantic Ocean and was sampled in the freshwater part of the lake (Suppl. Table 1).

Temperature, pH, conductivity and Secchi disk readings were measured *in situ*. In addition, surface water samples were collected in 10 l polypropylene containers for analyses of chemical and physical parameters. Technical details of the abiotic variables and chlorophyll-a measurements are described in Berman *et al.* (2018) and are summarized in Suppl. Table 1.

For filtered total plankton (FTP) samples, 50 ml of water from each lake was *in situ* filtered through a 50 µm nylon mesh to remove invertebrates, large protists and particles, and subsequently filtered through 0.22 µm pore-size polycarbonate filters (ø 47 mm; Millipore). The filters were flash frozen and kept at −80°C until DNA extraction. Genomic DNA from filters was extracted using a CTAB protocol (Fernández Zenoff, Siñeriz and Farías 2006).

### Flow cytometry cell sorting

Samples for flow cytometry (4.5 ml of water from each lake) were preserved with 0.5 ml of a solution containing glycerol and TE buffer, 1% final concentration (Rinke *et al.* 2014), deep frozen into liquid N_2_ and stored at −80°C. We used a FACSAria II flow cytometer cell-sorter (Becton-Dickinson) equipped with a blue laser (488 nm emission) and a red laser (633 nm). Fluorescent beads (1 μm, Fluoresbrite carboxylate yellow-green microspheres [Polysciences]) were added on each sample as internal standards. PPE were detected by their signature in plots of blue laser dependent red fluorescence of Chl-a (PerCP or FL3) versus size scatter (SSC-H) and discriminated from picocyanobacteria based on the red laser dependent red fluorescence (APC or FL4) of phycocyanin (Fig. 1). Between 50 000 and 100 000 PPE cells were sorted in “purity” mode into sterile cryotubes with 100 µl of buffer TE (Suppl. Table 2). Sterile PBS buffer was used as sheath fluid. Between 4 000 and 20 000 sorted cells were separated for epifluorescence microscopy observation (see below) and the rest were frozen (- 80°C) until DNA extraction.

**Fig. 1:**
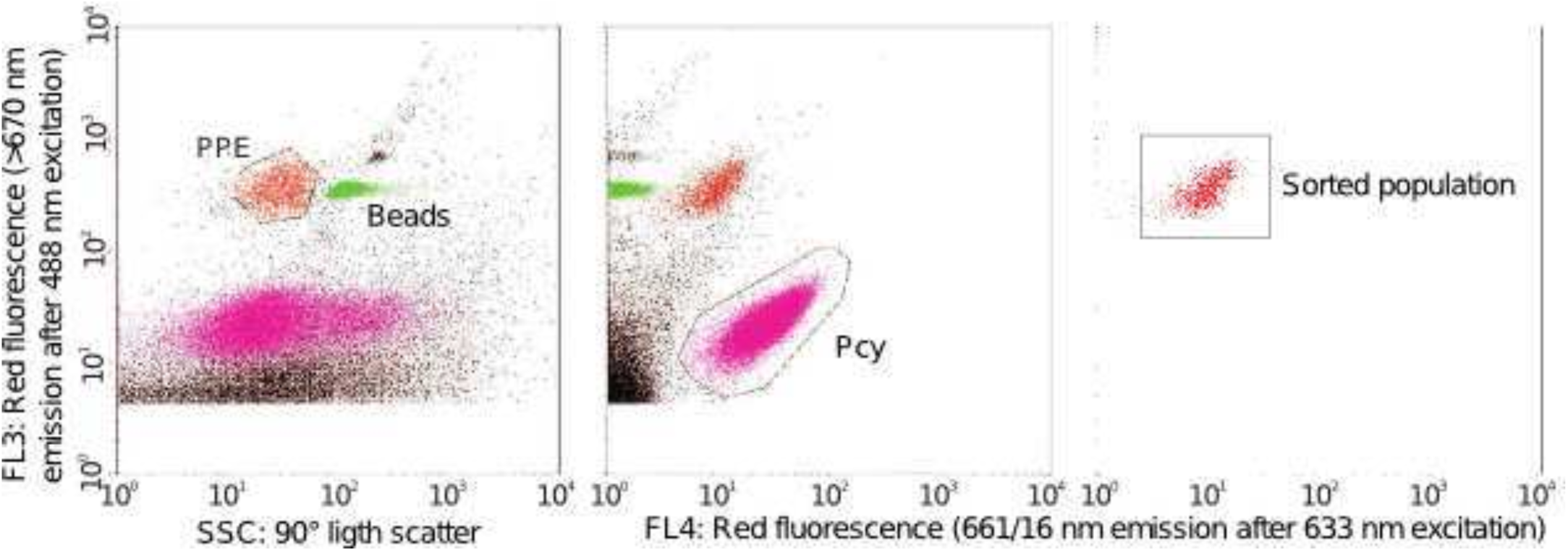
Example of a cytogram showing the strategy used to sort the photosynthetic picoeukaryotic (PPE) cells. PPE and Beads correspond to the cytometric populations of picoplankton and beads of 1 µm size, respectively.

### Epifluorescence microscopy

PPE sorted cells for epifluorescence microscopy were filtered through a 0.22 µm pore-size black polycarbonate filter and stained for counting following a standard DAPI staining procedure (Porter and Feig 1980). All filters were mounted onto a microscope slide with a drop of immersion oil for fluorescence (Immersol 518F) and stored at −20°C before observation. The relative abundance of distinct PPE morphotypes was quantified at 1 000x magnification using a Nikon Eclipse 80i microscope equipped with a HBO 50 W lamp and filter set for blue light excitation (BP 450–490 nm, FT 500 nm, LP 515 nm), green light excitation (BP 510–560 nm, FT 565 nm, LP 590 nm) and UV excitation (BP 340–380 nm, FT 400 nm, BP 435–485). Cell-size was estimated by image analysis using a colour camera (Nikon DS-Fi1) and Image-Pro Plus 4.5 software (Media Cybernetics). Between 26 and 50 cells were analysed for the most abundant morphotypes (i.e. spherical and spheroid) and at least 9 cells for less common ones.

### DNA extraction, PCR amplification and sequencing

Genomic DNA from sorted PPE was extracted adapting the CTAB protocol (Fernández Zenoff, Siñeriz and Farías 2006). In brief, cells were concentrated through centrifugation at 8000 rpm. CTAB lysis buffer, proteinase K (50 mM final concentration) and glycogen (5 mg mL^−1^) were added to the cells pellet according to Marie *et al.* (2010). Samples were then incubated at 56°C for 30 min and proteinase K inactivated at 75°C for 10 min. After, the protocol follows as described in Llames et al. (2013). For filtered total plankton sample (FTP) DNA extraction was performed using the CTAB protocol (Fernández Zenoff, Siñeriz and Farías 2006). Unfortunately, FTP sample from QU lake could not be sequenced due to technical problems in the sample preparation.

The V4 region of the 18S rRNA gene was amplified with nested and semi-nested PCR approaches for FTP and PPE sorted cells, respectively. The semi-nested PCR increased the sensitivity and yield which was not always the case with the nested protocol.

The first round of PCR amplification was performed with primers 63f and 1818r for FTP samples (Lepère *et al.* 2011) and V4f and 1818r for PPE samples (Piredda *et al.* 2017, Supp. Table 3). In both cases the thermal amplification conditions were: 95°C for 5 min, and 25 cycle of 98°C for 20 s, 52°C for 30 s, 72°C for 90s, followed by 5 min at 72°C for polymerase extension.

For the second round, the V4 fragment was amplified with specific Illumina primers V4F_illum (Stoeck *et al.* 2010) and V4R_illum (Piredda *et al.* 2017, Suppl. Table 3). Thermal conditions for FPT samples were: 95°C for 3 min, and 25 cycle of 98°C for 20 s, 65°C for 60 s, and 72°C for 90 s, followed by 5 min at 72°C for polymerase extension. For PPE samples, the thermal conditions applied were similar with a slight reduction of the annealing temperature and time to 52°C for 30 s. Final expected product length (ca. 406 bp) was checked by electrophoresis in 1% agarose gel. DNA bands were cut and purified with Nucleospin Gel and PCR Clean up (QIAGEN, USA). All amplifications were carried out in triplicate with Kapa HiFi polymerase and pooled before indexation with Nextera XT Kit. Negatives control used during PCR reactions were pooled and sequenced. The sequencing libraries and reagent preparation were performed according to Illumina manufacturer’s instruction (www.illumina.com/support) for MiSeq Reagent Kit V.3 for 600 cycles. Sequences have been deposited at the European Nucleotide Archive (ENA) under the BioProject number (PRJEB31039).

### Sequence processing

Reads were processed following the pipeline proposed by Logares (2017). Forward and reverse reads were corrected by SPAdes (Bankevich *et al.* 2012), the pair-merged products were performed by PEAR v0.9 (Zhang *et al.* 2014) and filtered using USEARCH v9 (Edgar 2010; Edgar 2013). OTUs were defined at 98% threshold of identity using command “uclust” from USEARCH v9 (Edgar 2010). Their taxonomic affiliation was assigned using BLAST (Altschul *et al.* 1997) and the curated database PR2 (gb203, Guillou *et al.* 2013). OTUs affiliated to Metazoa, Embryophyta, Archaea, Eubacteria and those that were relatively more abundant in negative controls respect to others samples were excluded from the analyses. Negative control sequences were probably produced during the PCR or were resulted of cross contamination during the sequencing step. Negative controls OTUs were affiliated mainly to Alveolates, Stramenopiles, Hacrobia and Archaeplastida. They had less than 4 reads in all samples and were present in less than 3 samples, including the negative control. Moreover, the contribution of these OTUs never exceeded 0.05% of the total reads per sample.

Sixteen OTUs, assigned to Basidiomycetes and Ascomycetes, were excluded from further analysis. The taxonomic identity of these OTUs revealed that they were all associated to human skin or to indoor environments and represented probably laboratory contaminants (Suppl. Table 4). They were completely absent in the corresponding FTP sample which suggest that they were accidentally incorporated during the cell sorting process.

OTUs represented by less than five reads were removed to avoid considering sequencing artefacts (Bokulich *et al.* 2012; Kircher, Sawyer and Meyer 2012; Nelson *et al.* 2014). The OTU affiliation was individually revised and those that had a percentage identity below 85% were Blasted against non-redundant (nr) database to find a closer match (Altschul *et al.* 1997).

OTUs were classified into pigmented and non-pigmented organisms based on their phylogenetic affiliation. OTUs affiliated to clades containing both types (i.e. Dinoflagellates, Chrysophyceae and Dictyochophyceae) were individually revised using NCBI database and functionally characterized based on their fine taxonomic affiliation (Suppl. Table 5).

### Data analysis

All data analyses were performed in R environment (http://cran.r-project.org). The relative abundance of OTUs among all samples (FTP and PPE) was compared after re-sampling of each sample at the minimum sample size (23 637 reads, from MA2 FTP sample) using the VEGAN package (Oksanen *et al.* 2017). In order to compare the richness between PPE and FTP, each pair of samples was re-sampled at the minimum number of reads from each site. The similarity between PPE samples was estimated using Bray-Curtis index after Hellinger transformation of data to normalize the OTU table (Legendre and Gallagher 2001). We used similar method described in del Campo and Massana (2011) to infer the novelty of an environmental sequence from PPE dataset. Briefly, we plotted the percentage of identity of each OTUs corresponding to the three principal classes of PPE found with the closest classified organism match (CCM) versus the closest uncultured/environmental match (CEM).

### Scanning electronic microscopy

Environmental samples preserved with 1% acidified Lugol’s iodine solution were filtered by 10 µm pore size mesh and washed several times with deionized water to remove the fixer. Subsequently, samples were oxidized with hydrogen peroxide (H_2_O_2_ 30%) and heating (60°C) for 24 h. Subsamples of the oxidized suspensions were washed thoroughly with deionized water, mounted in stubs and sputter-coated with a thin layer of gold using a combined metal / carbon deposition system (SPI SUPPLIES model SPI Module, cat. SPI 12157-AX) operated in Argon atmosphere. Samples were examined using a Phenom Pro scanning electron microscope at accelerating voltages of 5 and 10 kV, with a working distance of 2.5 ± 0.5 mm.

## RESULTS

### Sampling sites

The six selected lakes sampled are shallow (ca. 1.5 – 3 m depth), turbid (mean Secchi depth = 15 cm) and eutrophic (mean TP = 550 µg l^−1^) (Suppl. Table 1). Conductivity varied from 0.65 mS cm^−1^ in VI to 3.25 mS cm^−1^ in LJ, with the exception of QU that reached 12.46 mS cm^−1^. PPE abundance averaged 1.4 × 10^5^ cells ml^−1^ (Suppl. Table 2).

### Microscopy analysis of PPE

The sorted PPE cells size ranged from 1 to 3.3 µm (Suppl. Fig. 1a-b). Sorted cells were mostly grouped in two different morphotypes: spherical (average cell diameter: 2 µm; range: 1.0 - 3.3 µm) and the spheroid (average cell dimensions: 1.6 × 2.5 µm; range: 2.3 × 3.2 – 1.2 × 1.8 µm). These morphotypes accounted together for >80% of the cells in all the lakes (Table 1). No clear differences in cell-size were observed among lakes for any morphotype. In LI, we detected a large proportion of rectangular-shaped PPE (Suppl Fig. 1c), which accounted for about 16% of all cells (average cell dimensions: 2.2 × 1.2 µm; range: 2.5 × 1.4 – 1.8 × 0.9 µm), suggesting the presence of small pico-sized diatoms. Occasionally, other morphologies, i.e. flagellated, kidney-shaped and with thorns, were also detected. Only 2 out of 589 inspected cells were larger than the expected picoplanktonic size range. These cells belonged to the chlorophycean genera *Tetrastrum* and *Monoraphidium* respectively and were observed in CH sample.

**Table 1:**
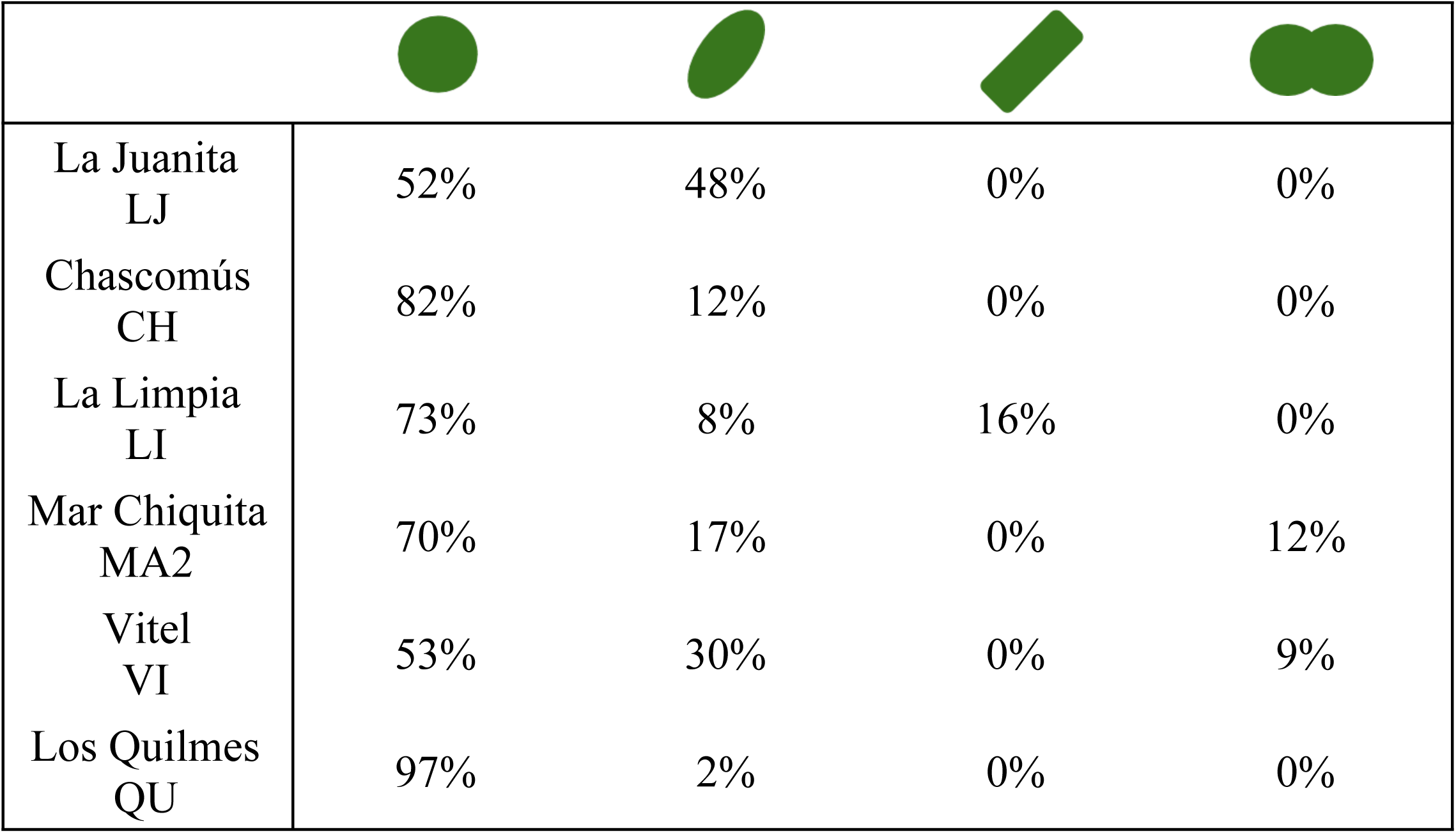
Percentage of morphotypes observed in each sample by fluorescence microscopy.

|  |  |  |  |  |
| --- | --- | --- | --- | --- |
| La Juanita<br>LJ | 52% | 48% | 0% | 0% |
| Chascomús<br>CH | 82% | 12% | 0% | 0% |
| La Limpia<br>LI | 73% | 8% | 16% | 0% |
| Mar Chiquita<br>MA2 | 70% | 17% | 0% | 12% |
| Vitel<br>VI | 53% | 30% | 0% | 9% |
| Los Quilmes<br>QU | 97% | 2% | 0% | 0% |

Scanning electronic microscopy (SEM) analyses performed on a sample from lake LI revealed the presence of *Aulacoseira granulata, A. granulata var. angustissima* and *A. ambigua*, with very thin cells of about 2.8-3.5 µm width (Suppl. Fig. 2). Some small specimens of *Nitzschia* sp. and *Pseudostaurosiropsis* sp. were also observed in SEM samples.

**Fig. 2:**
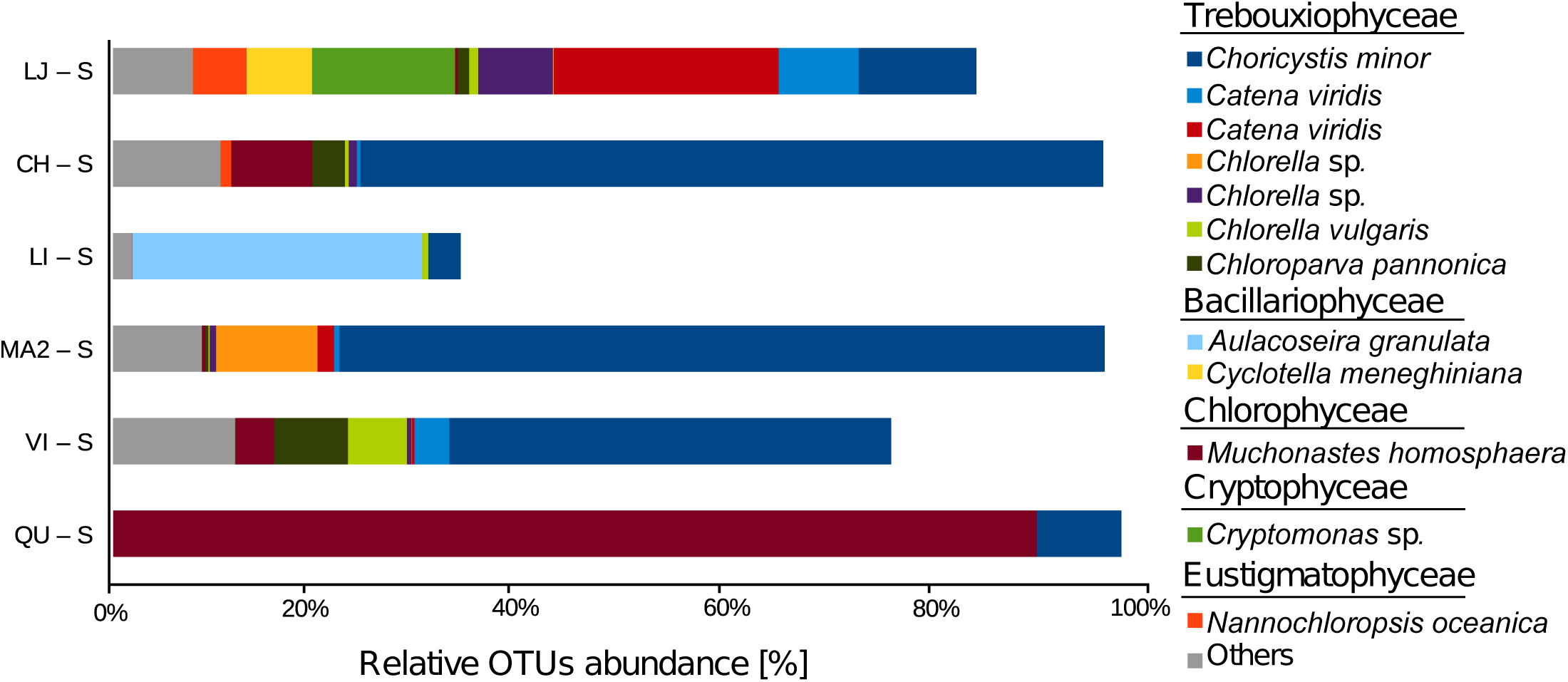
Relative abundance of dominant OTUs obtained from PPE samples (those that contributed with more than 1% in the total reads per sample). LJ (La J uanita); CH (Chascomús); LI (La Limpia); MA2 (Mar Chiquita); VI (Vitel); QU (Los Quilmes).

### Sequences analysis of sorted PPE

The number of reads retrieved per sample from PPE sorted cells ranged between 24 402 and 272 264 (Suppl. Table 6). Total of 257 OTUs were obtained with an average of 62 OTUs per lake (Suppl. Table 6). Among the 43 abundant OTUs (those that contributed with more than 1% in the total reads per sample), 24 were affiliated to Trebouxiophyceae, Chlorophyceae and Bacillariophyceae (Fig. 2, Suppl. Table 7). Overall, the PPE community composition among the lakes showed low similarity with Bray-Curtis dissimilarity index above 0.63 (Suppl. Table 8).

Around 81.7% of the reads could be affiliated to pigmented organisms. Trebouxiophyceae dominated PPE in LJ, CH, MA2 and VI, contributing with 50 to 91% of the sequences. Chlorophyceae and Bacillariophyceae dominated in the two other lakes, QU and LI, with 88 and 27% of the sequences respectively (Fig. 3). Other classes like Pedinophyceae, Eustigmatophyceae, Mammiellophyceae, Cryptophyceae and Chrysophyceae were less represented.

**Fig. 3:**
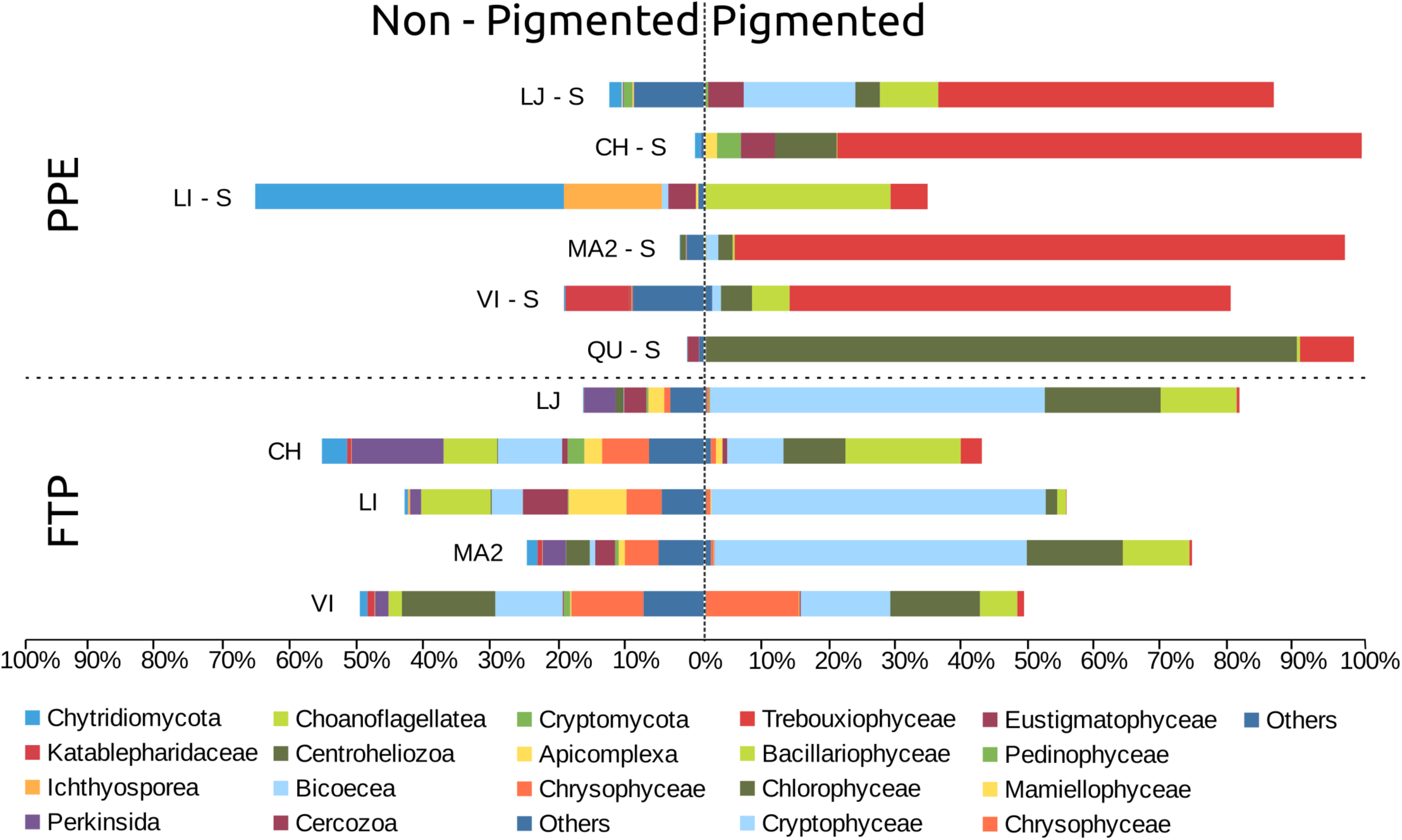
Relative abundances of the main classes for each sample of PPE sorted cells and FTP (filtered total plankton). Organisms were discriminated between pigmented and non-pigmented. LJ (La J uanita); CH (Chascomús); LI (La Limpia); MA2 (Mar Chiquita); VI (Vitel); QU (Los Quilmes).

Trebouxiophyceae OTUs were represented mostly by genera *Choricystis, Chlorella* (*sensu stricto*), *Catena* and *Nannochloris.* Only OTU_1 related to *Choricystis minor* (>99.5% of identity) was present in all lakes (Fig. 2, Suppl. Table 7). Chlorophyceae were mainly represented by OTU_4, which was affiliated to the genus *Mychonastes* (>99.2% of identity). This single OTU accounted for more than 88% of the sequences in lake QU. Bacillariophyceae were represented by 3 OTUs affiliated to *Aulacoseira* and *Cyclotella* genera, with more than 99.0% and 98.5% of identity respectively. Less abundant groups of PPE were represented by OTUs related to *Monodus subterranea* and *Nannochloropsis oceanica* (Eustigmatophyceae), *Chlorochytridion* spp. (Pedinophyceae), *Cryptomonas* spp. (Cryptophyceae), *Crustomastix* and *Monomastix* (Mamiellophyceae). Only 9 out of the 43 abundant OTUs were present in at least two samples with high abundance. Most of them corresponded to photosynthetic organisms, except for OTU_152 that was a Rhizarian and thus probably a heterotroph (Suppl. Table 7).

Non-pigmented groups and unknown according to our functional classification accounted for 18.1% and 0.2% of the reads respectively (Fig. 3, Suppl. Table 5). Chytridiomycetes and Ichthyosporea, two groups that include diatom parasitoids, were particularly abundant in the lake dominated by diatoms (LI).

In order to investigate the degree of genetic novelty of the OTUs retrieved from the PPE sorted cells from the three dominant classes (Trebouxiophyceae, Chlorophyceae and Bacillariophyceae), we plotted the percentage of identity of each OTUs with the closest classified organism match (CCM) versus the closest uncultured/environmental match (CEM). About 22% of all OTUs (i.e. 12% of Trebouxiophyceae, 7% of Chlorophyceae and 3% of Bacillariophyceae) had less than 95% of identity with any sequence reported (Fig. 4).

**Fig. 4:**
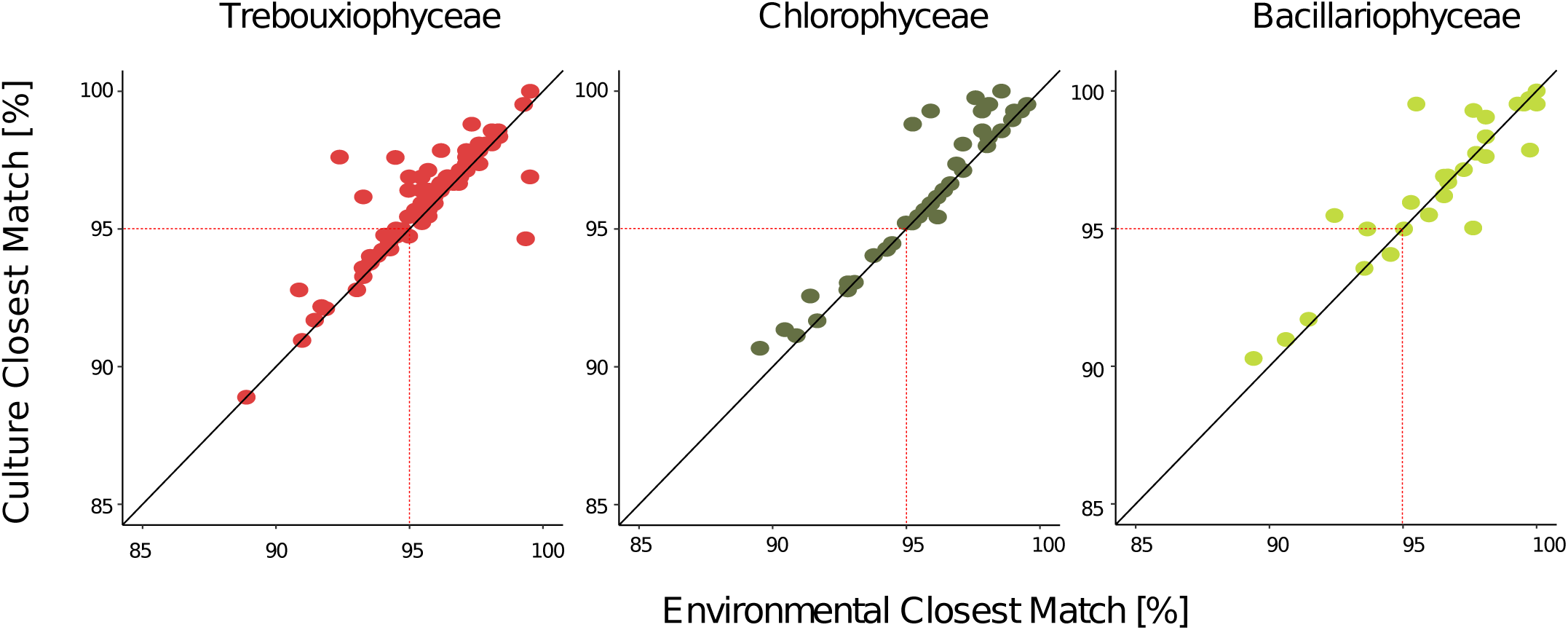
Novelty pattern derived from OTUs among the three main PPE classes. Dots represent the % similarity with the closest environmental match (CEM) and the closest cultured match (CCM) for each sequence within the three taxa (Trebouxiophyceae, Chlorophyceae and Bacillariophyceae sequences, respectively).

### Comparison between PPE and FTP samples

The number of reads per sample retrieved from the FTP varied in a range between 23 637 – 275 343. After samples standardization, 68% of the reads were affiliated to known pigmented organisms, mainly Cryptophyceae (33.9%), Chlorophyceae (11.3%), Bacillariophyceae (9%) and Chrysophyceae (3.2%). Trebouxiophyceae accounted only for 1% of the reads (Fig. 3). Non-pigmented groups (39% of the reads) were very diverse with Perkinsida, Chytridiomycota, Apicomplexa, Centroheliozoa and Bicosoecida being the most abundant.

A total of 916 OTUs were identified in FTP samples, with an average of 314 OTUs per lake and only 123 OTUs (ca. 13%) shared between PPE sorted samples. Among them, 71 were affiliated to pigmented group and 52 to non-pigmented.

Among pigmented groups, the richness per lake was in general higher in FTP as compared to sorted samples (Fig. 5), except for Trebouxiophyceae that was three times richer in sorted samples (92 OTUs from sorted samples versus only 30 in FTP). Similar trend was observed for Pedinophyceae in CH and Eustigmatophyceae in CH and LJ. In contrast, the richness of Chlorophyceae and Bacillariophyceae were higher in FTP than PPE (Fig. 5). Very few OTUs affiliated to these classes were shared between PPE and FTP (Fig. 6). Likewise, the affiliation of dominant OTUs for each algal class in FTP differed from the PPE samples. Trebouxiophyceae was dominated by *Micractinium pusillum, Chlamydomonas spp.* dominated Chlorophyceae and *Cyclotella, Chaetoceros* and *Amphora* were the most abundant Bacillariophyceae. These organisms have larger cells size, well above pico-size.

**Fig. 5:**
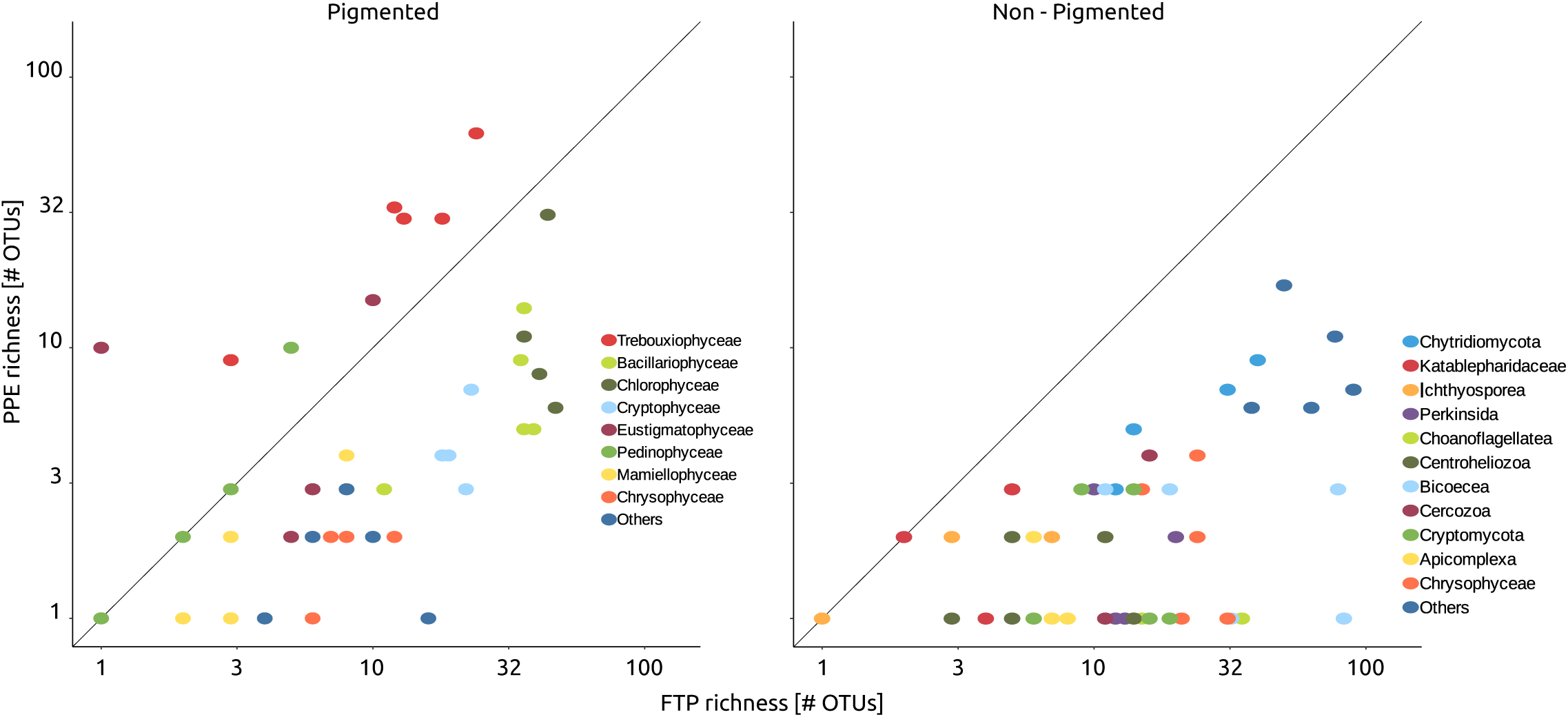
Richness (number of OTUs) of sorted samples (i.e. PPE) against richness from filtered plankton (FP) sample, discriminated by groups.

**Fig. 6:**
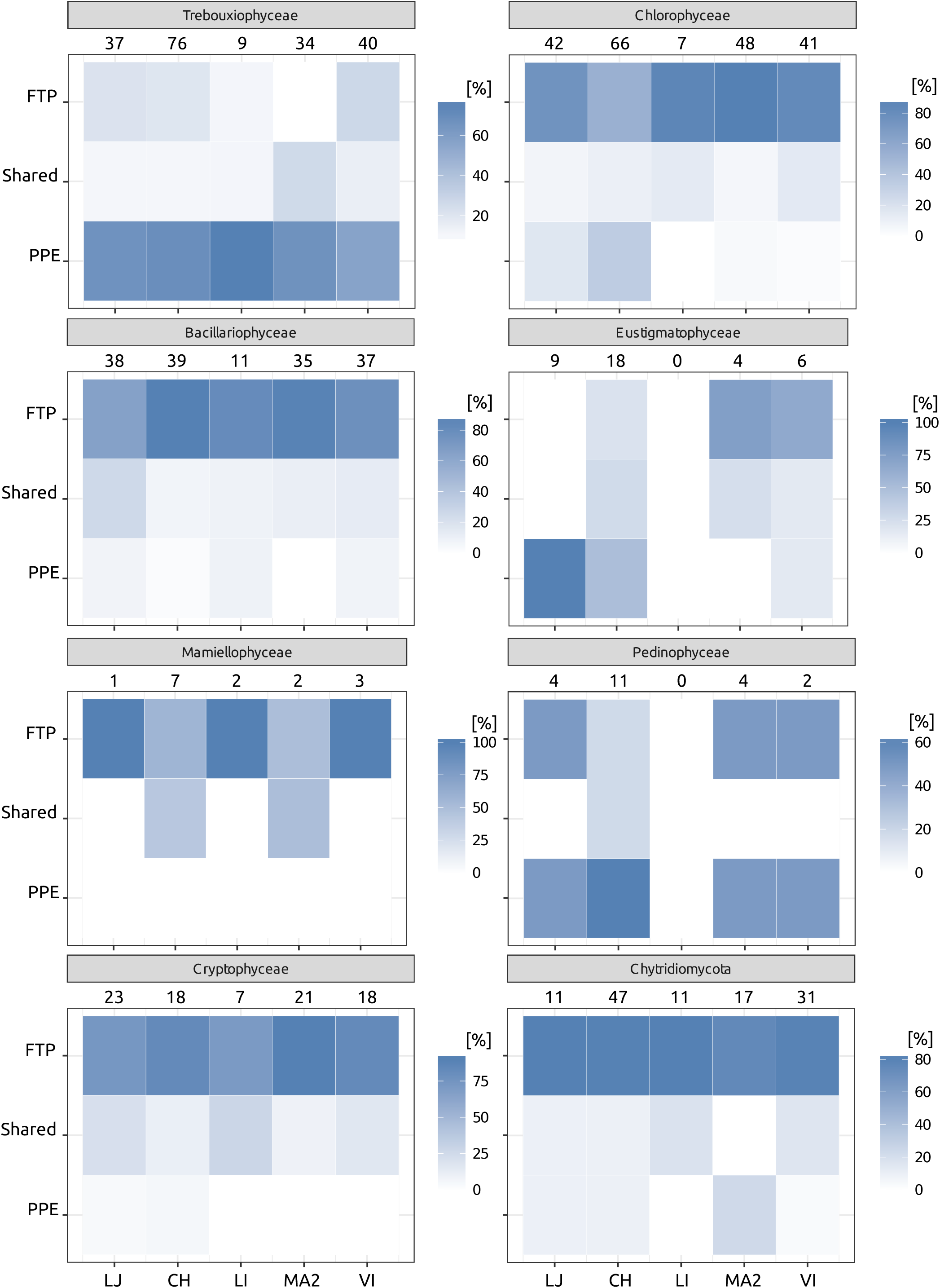
Heatmaps showing the proportion of OTUs for the main clades, that appeared exclusively in the sorted samples (PPE) or in the filtered total plankton (FTP) sample, and those that were sheared. Numbers indicate the total amount of OTUs for each clade and lake.

## DISCUSSION

Several authors suggested sequencing of sorted cells as a suitable alternative to overcome some biases, like the fact that picoplanktonic photosynthetic eukaryotes are outnumbered by heterotrophic cells (Shi *et al.* 2009; Marie *et al.* 2010; Balzano *et al.* 2012; Li *et al.* 2017) or the sequential filtering that still may allow large non-picoplanktonic organisms to pass through the filters in the picoplanktonic fraction. This problem is particularly critical in highly turbid environments like eutrophic shallow lakes.

We observed marked differences in community composition between FTP and PPE cell sorting samples. Yet, a total of 162 OTUs were common between cell sorting and FTP samples. PPE OTUs assigned to Trebouxiophyceae dominated sorted samples (50% of total reads) while FTP contained mostly large sized (over 10 µm) phototrophs (i.e. Cryptophyceae, diatoms among 34% and 9% of the total reads, respectively) or heterotrophic organisms, such as Perkinsida (5% of total reads) and Bicoecea (5% of total reads). The class Trebouxiophyceae, that contains photosynthetic eukaryotes for which most members are smaller than 3 µm in size were poorly represented in FTP. Trebouxiophyceae were three times richer in cell sorting samples (92 OTUs) than FTP (30 OTUs) illustrating that even large sequencing effort as provide by Illumina MiSeq fails to reveal a more complete picture of PPE diversity filtered samples, even when rarefaction curves approached asymptote (Suppl. Fig. 3).

Cell-sorting methodology have been applied by several studies aiming to unveil PPE diversity from marine samples (Shi *et al.* 2009; Marie *et al.* 2010; Balzano *et al.* 2012). However, similar approach to freshwater samples have been applied only twice (Li *et al.* 2017; Shi *et al.* 2018) to our knowledge. These authors observed the presence of dominant OTUs (e.g. *Nitzschia palea, Chlamydomonas* spp. and *Ankyra lanceolata*) corresponding to organisms measuring over 10 µm in length (Krientz and Heynig 1982; Pröschold *et al.* 2001; Trobajo *et al.* 2009), which clearly do not belong to the PPE fraction. This highlights the importance of a right gating strategy using for cell sorting (Suppl. Fig. 3 from Li et al. 2017) and the needs for establishing controls. These authors firstly selected the smallest cells based on their low FSC signal, and then separate eukaryotic phototrophs from cyanobacteria in a secondary FL3 (chlorophyll) vs FL5 (phycocyanin) plot. By using this approach, they failed to discriminate pico-from nano-sized phototrophic eukaryotes, thus pooling both size classes in the same gate. We added 1 µm latex beads to our samples as an internal standard for size and used a different gating strategy (Fig. 1), very similar to the one used by Marie *et al.* (2010). Using this approach PPE always falls to the left side of the beads (with less SSC) usually forming a single population with similar or slightly higher red fluorescence signal in the FL3-SSC plot. Our results were validated with epifluorescence microscopy, which corroborated that 99% of the sorted cells had the expected PPE size (1 – 3 µm).

Comparisons with other studies that used the same methodology (i.e. sorting and MiSeq) are scarce, precluding a definitive conclusion about the importance of the diversity of PPE from these shallow lakes in relation to other environments. For instance, the two studies performed in eutrophic Chinese shallow lakes observed between 51 and 367 OTUs per sample (Li *et al.* 2017; Shi *et al.* 2018), while here we obtained between 27 and 141 OTUs. However, as mentioned above, the gating strategy used in Li *et al.* (2017) and Shi *et al.* (2018) included also nanoplanktonic algae. In marine samples, the combination of cell sorting and sequencing to specifically study PPE was applied several times (e.g. Shi *et al.* 2009; Marie *et al.* 2010), though only in one occasion high throughput sequencing was applied (Ribeiro *et al.* 2017). In this case, the total richness found in pico- and nano-size fraction together was 258 OTUs, the same richness obtained in the present work only in the picoplanktonic fraction.

OTUs assigned to Cryptophyceae (i.e. *Cryptomonas*) were present in the PPE sorted samples (Fig. 3) despite the fact that our settings discriminated against phycoerythrin-containing cells. These OTUs accounted for ∼10% of the reads from LJ sorted sample and same OTUs also dominate FTP sample (Suppl. Table 6). The presence of phycobilin-containing algae in the sorted population could be explained by a cross-contamination between the pico- and nano-sized populations. However, it is worth mentioning that cryptophytes could not be distinguished in this sample when observed under epifluorescence, even though they are easy to identify under blue and green light excitation. Another plausible explanation could be the low phycobilin content of individual cells of pico-sized cryptophyte.

We detected a significant amount of sequences (18.1% of total reads) belonging to non-pigmented organisms from sorted PPE samples. Besides the likely contaminating organisms already present in the cell sorter (i.e. fungi, Suppl. Table 3) most of these sequences were affiliated to parasites or predators of PPE. Most non-pigmented OTUs were affiliated to known freshwater algal parasites like Chytridiomycota and Cryptomycota (Lara, Moreira and López-García 2010; Jones *et al.* 2011; Sime-Ngando, Lefèvre and Gleason 2011; Rasconi, Niquil and Sime-Ngando 2012), and the obligate parasitic group Ichthyosporea (Glockling, Marshall and Gleason 2013). Chytridiomycota (OTU_17) and Ichthyosporea (OTU_29) were particularly abundant in lake LI where PPE was dominated by the diatom from genus *Aulacoseira* (OTU_24). This genus is known to act as a host for several Chytridiomycota, including the genus *Rhizophydium* OTU_54 (Canter 1967; Seto, Kagami and Degawa 2017). Ichthyosporea is a very diverse clade that includes animal symbionts (both parasites and mutualists). To date, there is no formal proof that they can be associated to pico-sized algae, but two environmental clades, one of them that include OTU_29, lack cultured representatives and are often found in environmental DNA-based diversity surveys of freshwater plankton (Reynolds et al., 2017). This suggests that these organisms are either pigmented (which is highly unlikely given the parasitic nature of the whole group) or are associated with PPE. Our results illustrate how little is known about host – parasite interaction in freshwater PPE communities. Furthermore, a Rhizarian OTU (number 152) was also encountered abundantly in QU lake (1.7% of reads), probably representing a PPE predator or a parasite since specialized parasites of algae have been well characterized within Rhizaria (Hess and Melkonian 2013). None of the above-mentioned OTUs were abundant enough to be detected in FTP samples, suggesting that they are indeed associated to sorted PPE.

Microscopy revealed the presence of two different main morphotypes of PPE, spherical and spheroid, which accounted for >80% of the cells in all samples. The spherical morphotype for example, dominated in both QU and CH (representing, respectively, 97 and 82% of all PPE), however, while the most abundant OTU in the PPE community from QU (OTU_4) was affiliated to *Mychonastes homosphaera* (>99.2% Identity; Chlorophyceae), CH was dominated by OTU_14, closely related to *Choricystis minor* (>99.5% Identity; Trebouxiophyceae). This illustrates a long-standing problem, the lack of easily distinguishable features in most PPE species that prevents morphology-based diversity surveys for small-size plankton.

A rectangular shape morphotype (Suppl. Fig. 1c) was only found in the PPE of lake LI, dominated by diatoms. This cell-shape could be potentially related to some picosized diatom species. The existence of minute diatoms was previously reported, and cultured, in marine and freshwater systems (Collier and Murphy 1962; Sar et al. 2012). In lake LI, sequences of pigmented PPE were dominated by OTU_24 assigned to *Aulacoseira granulata* (99.1% Identity). The phytoplankton estimated using standard inverted microscope methodology showed that the genus *Aulacoseira* accounted for 88% of the total abundance (Izaguirre, unpublished data; Izaguirre *et al.* 2014), while scanning electronic microscopy revealed the presence few specimens of small *Nitzschia* sp. and *Pseudostaurosiropsis* sp., and many cells of *Aulacoseira* represented by three different species: *A. granulata* var. *granulata, A. granulata* var. *angustissima* and *A. ambigua* (Suppl. Fig. 2). The later species has very thin cells that measured about 2.8-3.5 µm in diameter, although with a mean mantle height of 15 µm. O’Farrell et al (2001) analyzed morphological variability of *Aulacoseira granulata* in the Lower Paraná River and found that mean length was inversely correlated with suspended solids content. The authors reported minimum cell length of 7 µm and cell diameter of 2 µm for specimens analyzed. Lake LI is characterized by turbid conditions that would favor the dominance of small specimens of *Aulacoseira* spp. The dominance of sequences of *Aulacoseira* among the PPE sorted can also be explained by the existence of individuals with shorter cells than those found it in SEM, probably as a result of being actively dividing. Finally, we cannot discard the possibility that some broken cells have fallen within the population of PPE population in the FL3-SSC plot (Fig. 1).

Even though the lakes sampled shared many physical-chemical features and are placed on the same geographic and climatic region, the similarity of PPE composition among the samples was low with respect to the high Bray-Curtis values calculated (Suppl. Table 8). The two most striking differences were observed in QU and LI. The former was dominated by a single OTU belonging to chlorophyceaen *Mychonastes homosphaera* and the latter by diatoms. The high conductivity recorded in QU might explain the dominance of *Mychonastes homosphaera*, while the predominance of inorganic turbidity in LI, instead of organic turbidity that predominate in Pampean lakes (Fermani *et al.* 2014), might have favoured the dominance of diatoms that are known to be adapted to growth in turbid systems (Allende *et al.* 2009). The other four lakes were clearly dominated by Trebouxiophyceae, the richest algal class among the PPE. Similar predominance was reported during winter in other eutrophic shallow lakes from the northern hemisphere (Felföldi *et al.* 2011; Li *et al.* 2017). The cosmopolitan *Choricystis minor* was the only OTU that appear in all lakes and was always highly abundant. In contrast, the rest of the abundant OTUs were present in, at most, one or two lakes. This high variation between lakes community compositions cannot be explained by the low connectivity among lakes, since very close lakes, like LI, CH and VI, harbour very different PPE composition. Moreover, CH and VI are directly connected by a short stream, suggesting that local forces would be important drivers of the community structure of PPE, as it was already determined for heterotrophic bacteria in three lakes from the same area (Llames *et al.* 2017).

In marine systems, PPEs are mainly represented by Mamiellophyceae and Chloropicophyceae (Prasinophyta), Prymnesiophyceae, Chrysophyceae, among others (Jardillier *et al.* 2010; Worden *et al.* 2012; Gómez-Pereira *et al.* 2013; Unrein *et al.* 2014; Lopes dos Santos *et al.* 2017). In particular, Prasinophytes that are widespread in oceans and usually dominate PPE (Vargas et al. 2015; Vannier et al. 2016) were nearly absent in our samples. In contrast to marine plankton, Trebouxiophyceae and Chlorophyceae dominate PPE communities in shallow lakes, which are probably likely the ecological counterparts of marine PPE Prasinophyceae. The few available results suggest that the composition of PPE in freshwater clearly differs from marine systems.

Interestingly, we found a high percentage of novelty in our cell-sorted samples, reinforcing that this approach is suitable for new taxa discovery. Indeed, more than 22% of OTUs affiliated to Trebouxiophyceae, Chlorophyceae and Bacillariophyceae had less than 95% of identity with any sequence from public databases. This is particularly remarkable since these organisms, usually easy to culture, are represented by many voucher specimens in culture collections, and have been intensively bar-coded (Krienitz, Huss and Bock 2015).

Our study showed that freshwater planktonic diversity remains largely to be explored, especially in understudied parts of the world such as the Pampean Lakes in Central Argentina, and the power of combining cell sorting and high throughput sequencing in limnology studies of PPE community composition as well as to describe novel diversity.

## ACKNOWLEDGEMENTS

We gratefully thank Roberto Escaray, José F. Bustingorry and Nadia Diovisalvi for the technical assistance in lake sampling and chemical analyses. We also thank Horacio Zagarese for helpful comments on the manuscript. We are most grateful to the Biogenouest Genomics Genomer platform core facility for its technical support.

## FUNDING

This study was supported by the National Geographic Society grant #9736-15 and the ANPCyT “Agencia Nacional de Promoción Científica y Tecnológica” (PICT-2014-1290; PICT-2016-1079). EL was funded by the program “Atracción de talentos CAM” 2017-T1/AMB-5210.

