## Supplementary Material for "Diversity of photosynthetic picoeukaryotes in eutrophic shallow lakes as assessed by combining flow cytometry cell-sorting and high throughput sequencing"

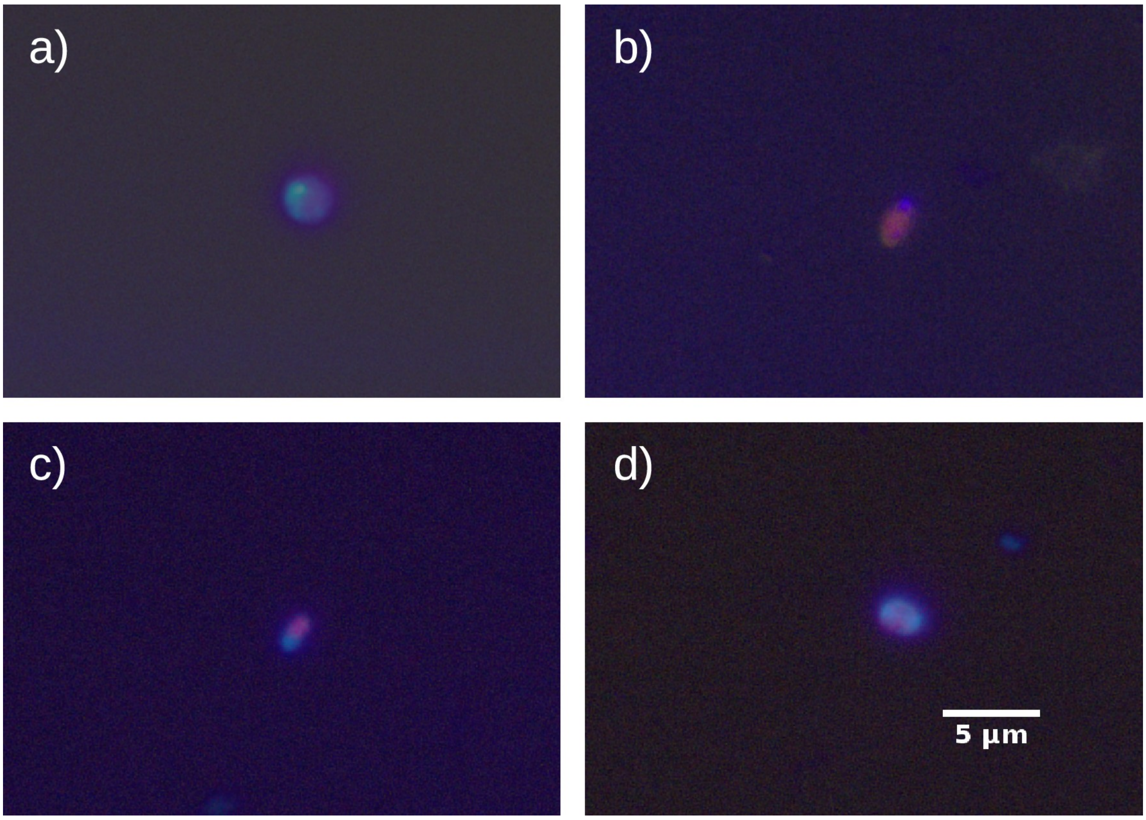


Supplementary Fig. 1: Epifluorescence micrography of most abundant PPE morphotypes: (a) spherical; (b) spheroid; (c) rectangular-shaped and (d) cells that are apparently in-division with two nucleus-like particles. Pictures after excitation with UV radiation showing the blue nucleus stained with DAPI and pictures after blue light excitation showing red chloroplasts were merged in only one picture.

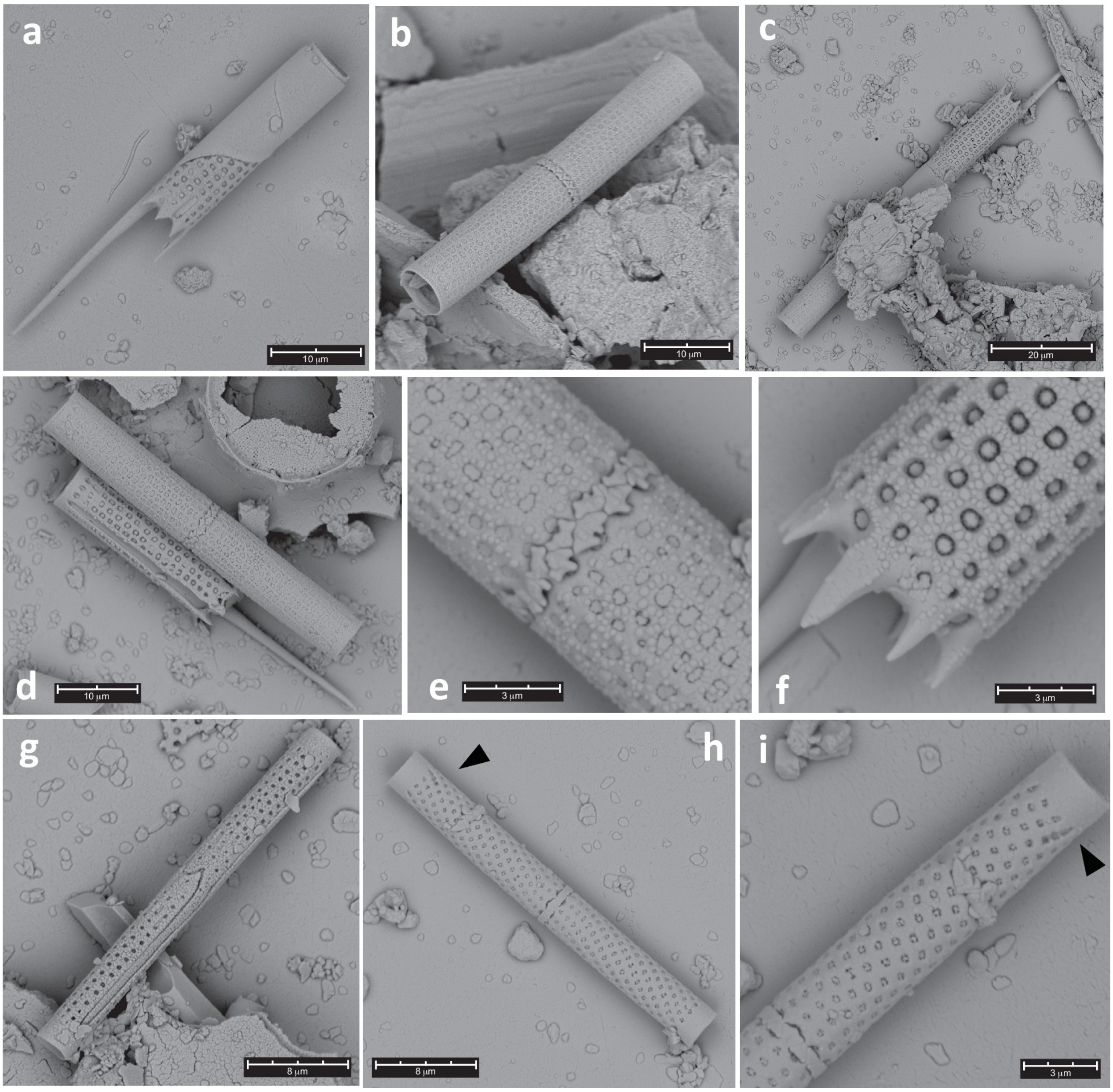


Supplementary. Fig. 2: Electron micrographs of the external surface of *Aulacoseira* valves (SEM). a-f) *Aulacoseira granulata* (Ehrenberg) Simonsen, g) *A. granulata* var. *angustissima* (Müller) Simonsen, h-i) *A. ambigua* (Grunow) Simonsen thin specimens, external opennings of the elongate rimoportula (arrows).

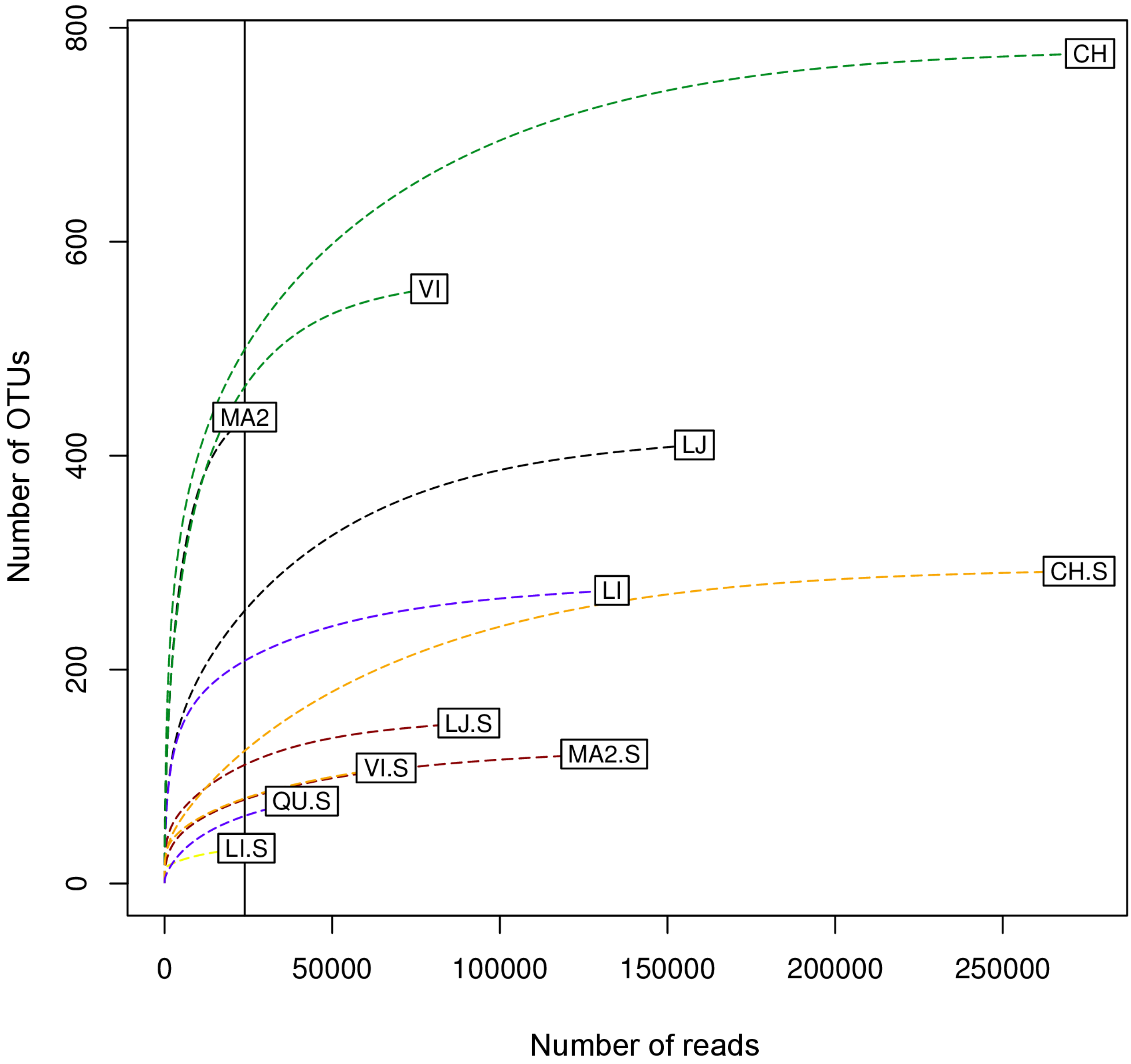


Supplementary Fig 3: Rarefaction curves of V4 SSU rRNA gene sequences in each sample before normalization of the minimal number of sequences (23902).

Supplementary Table 1: Morphometrical and physico-chemical features of the six lakes studied.

| | La Juanita (LJ) | Chascomús (CH) | La límpia (LI) | Mar Chiquita (MA2) | Vitel (VI) | Los Quilmes (QU) |
|---|---|---|---|---|---|---|
| Sampling day | 30/09/15 | 17/09/15 | 14/09/15 | 15/09/15 | 14/10/15 | 01/12/15 |
| Latitude | -38.522 | -35.582 | -35.602 | -37.650 | -35.545 | -35.881 |
| Longitude | -60.140 | -58.042 | -57.812 | -57.406 | -58.112 | -62.873 |
| Area ($km^2$) | 1 | 31 | 6 | 45 | 15 | 13 |
| Perimeter (km) | 5.33 | 32.34 | 14.07 | 69.07 | 22.62 | 18.76 |
| T (°C) | 10 | 15 | 15 | 18 | 17 | 26 |
| Secchi depth (cm) | 9 | 10 | 14 | 10 | 7 | 40 |
| pH | 8.44 | 8.14 | 8.78 | 8.56 | 8.94 | 8.53 |
| Conductivity (mS $cm^{-1}$) | 3.25 | 0.79 | 1.14 | 2.64 | 0.64 | 12.46 |
| NK (mg $l^{-1}$) | 7.47 | 5.05 | 4.76 | 4.86 | 4.64 | 6.72 |
| TP (mg $l^{-1}$) | 0.73 | 0.55 | 0.84 | 0.25 | 0.44 | 0.49 |
| Chl-a (µg $l^{-1}$) | 475 | 51 | 10 | 53 | 88 | 17 |

T, water temperature; NK, total Kjeldahl nitrogen; TP, total phosphorus and Chl-a, phyoplanktonic chlorophyll-a

Supplementary Table 2: Abundance and number of PPE sorted cells.

| Lake name | Sample ID | PPE abundance [cell ml-¹] | # of sorted cells for sequencing | # of sorted cells for microscophy |
|---|---|---|---|---|
| La Juanita | LJ | 326 000 | 80 000 | 20 000 |
| Chascomús | CH | 80 300 | 45 000 | 5 000 |
| La Limpia | LI | 65 800 | 87 800 | 15 000 |
| Mar Chiquita | MA2 | 59 700 | 103 000 | 10 000 |
| Vitel | VI | 113 000 | 104 686 | 4 000 |
| Los Quilmes | QU | 202 000 | 80 000 | 20 000 |

Supplementary Table 3: Primers sequences and references.

| Target | Primer | Illumina Tail | Sequence (5’ - 3') | Reference |
|---|---|---|---|---|
| 18S rRNA | 63F | - | ACG CTT GTC TCA AAG ATT A | Lepère *et al*. 2011 |
| 18S rRNA | 1818R | - | ACG GAA ACC TTG TTA CGA | Lepère *et al*. 2011 |
| 18S rRNA (V4) | V4F_illum | TCG TCG GCAG CGT CAG ATG TGT ATA AGA GAC AG | CCA GCA SCY GCG GTA ATT CC | Piredda *et al*. 2017 |
| 18S rRNA (V4) | V4R_illum | GTC TCG TGG GCT CGG AGA TGT GTA TAA GAG ACA G | ACT TTC GTT CTT GAT YRA TGA | Piredda *et al*. 2017 |

Supplementary Table 4: Contaminant OTUs affiliated to Basidiomycota and Ascomycota.

| | Sorting Samples (PPE) | | | | | | Environmental Samples (FTP) | | | | | OTU related affiliation after revision | | | | |
|---|---|---|---|---|---|---|---|---|---|---|---|---|---|---|---|---|
| OTU | LJ - S | CH - S | LI - S | MA2 - S | VI - S | QU - S | LJ | CH | LI | MA2 | VI | % of Identity | NCBI ACCN | Affiliation | Bibliography description | Reference |
| OTU_43 | 136 | 1147 | 1091 | 30 | 77 | 3693 | 0 | 3 | 9 | 3 | 2 | 99,75 | EU192367 | *Malassezia restricta* | Often skin associated. | Prohic, A., Simic, D., Sadikovic, T. J., & Krupalija-Fazlic, M. (2014). Distribution of Malassezia species on healthy human skin in Bosnia and Herzegovina: correlation with body part, age and gender. Iranian Journal of Microbiology, 6(4), 253–262. |
| OTU_80 | 25 | 243 | 1784 | 0 | 11 | 152 | 0 | 0 | 0 | 0 | 0 | 99,28 | KX951409 | *Epicoccum nigrum* | Highly robust and ubiquitous fungus, frequent in indoor habitats. | Singh, edited by Jagjit (1994). Building Mycology Management of Decay and Health in Buildings. (1st ed.). Hoboken: Taylor & Francis Ltd. ISBN 0-203-97473-5. |
| OTU_85 | 17 | 166 | 1474 | 2 | 0 | 699 | 0 | 0 | 0 | 0 | 1 | 99,28 | AB214655 | *Verticillium insectorum* | Also found in indoor habitats. | Sundh, I., (2012) Beneficial Microorganisms in Agriculture, Food and the Environment: Safety Assessment and Regulation. CAB International. ISBN 9781780640082 |
| OTU_21 | 2 | 1 | 8121 | 138 | 3 | 4 | 0 | 0 | 0 | 0 | 3 | 99,52 | KC674536 | Uncultured fungus | Isolated from skin, plantar hell. Host = "Homo sapiens". | Findley, K., Oh, J., Yang, J., Conlan, S., Deming, C., Meyer, J. A., … Segre, J. A. (2013). Human Skin Fungal Diversity. Nature, 498(7454), 367–370. http://doi.org/10.1038/nature12171 |
| OTU_226 | 1 | 0 | 523 | 0 | 0 | 0 | 0 | 0 | 0 | 0 | 0 | 99,76 | KU167832 | *Rhodotorula mucilaginosa* | Also often found indoor or in clinical samples. | Arendrup, M. C., Boekhout, T., Akova, M., Meis, J. F., Cornely, O. A., & Lortholary, O. (2014). ESCMID and ECMM joint clinical guidelines for the diagnosis and management of rare invasive yeast infections. Clinical Microbiology and Infection, 20(s3), 76-98. |
| OTU_10062 | 0 | 4 | 0 | 0 | 0 | 0 | 0 | 0 | 0 | 0 | 0 | 89,56 | JF784006 | *Demodex folliculorum* | Host="Homo sapiens". | Kligman, A. M., & Christensen, M. S. (2011). Demodex folliculorum: requirements for understanding its role in human skin disease. The Journal of investigative dermatology, 131(1), 8. |
| OTU_1103 | 0 | 0 | 4 | 0 | 0 | 0 | 0 | 0 | 0 | 0 | 0 | 93,54 | HQ231228 | *Rhodotorula sp.* | Also often found indoor or in clinical samples. | Arendrup, M. C., Boekhout, T., Akova, M., Meis, J. F., Cornely, O. A., & Lortholary, O. (2014). ESCMID and ECMM joint clinical guidelines for the diagnosis and management of rare invasive yeast infections. Clinical Microbiology and Infection, 20(s3), 76-98. |
| OTU_1252 | 0 | 0 | 0 | 50 | 0 | 0 | 0 | 0 | 0 | 0 | 0 | 96,39 | KC674536 | Uncultured fungus | Isolated from skin, plantar hell. Host = "Homo sapiens". | Findley, K., Oh, J., Yang, J., Conlan, S., Deming, C., Meyer, J. A., … Segre, J. A. (2013). Human Skin Fungal Diversity. Nature, 498(7454), 367–370. http://doi.org/10.1038/nature12171 |
| OTU_1288 | 0 | 0 | 4 | 0 | 0 | 0 | 0 | 0 | 0 | 0 | 0 | 97,86 | KU317623 | *Rhodotorula mucilaginosa* | Also often found indoor or in clinical samples. | Arendrup, M. C., Boekhout, T., Akova, M., Meis, J. F., Cornely, O. A., & Lortholary, O. (2014). ESCMID and ECMM joint clinical guidelines for the diagnosis and management of rare invasive yeast infections. Clinical Microbiology and Infection, 20(s3), 76-98. |
| OTU_160 | 0 | 0 | 1165 | 0 | 1 | 0 | 0 | 0 | 0 | 0 | 0 | 99,52 | KF036672 | *Dioszegia takashimae* | Basidiomycetous yeast; air sample; contaminated aquifer sediment; sulfide-rich Zodletone spring; plant leaves | NCBI: Related HITs source |
| OTU_23 | 0 | 2 | 2464 | 10 | 1 | 10194 | 0 | 0 | 0 | 1 | 0 | 99,52 | KJ708442 | *Sporobolomyces marcillae* | Found in indoor habitats. | Ronald E. Gots , Nancy J. Layton & Suellen W. Pirages (2003): Indoor Health: Background Levels of Fungi, AIHA Journal, 64:4, 427-438. |
| OTU_260 | 0 | 141 | 322 | 0 | 0 | 310 | 0 | 7 | 0 | 0 | 0 | 99,52 | KJ701198 | *Sporobolomyces oryzicola* | Found in indoor habitats. | Ronald E. Gots , Nancy J. Layton & Suellen W. Pirages (2003): Indoor Health: Background Levels of Fungi, AIHA Journal, 64:4, 427-438. |
| OTU_3583 | 0 | 1 | 0 | 0 | 0 | 26 | 0 | 0 | 0 | 0 | 0 | 97,04 | KC675128 | Uncultured fungus | Isolated from skin, plantar hell. Host = "Homo sapiens". | Findley, K., Oh, J., Yang, J., Conlan, S., Deming, C., Meyer, J. A., … Segre, J. A. (2013). Human Skin Fungal Diversity. Nature, 498(7454), 367–370. http://doi.org/10.1038/nature12171 |
| OTU_711 | 0 | 0 | 4 | 0 | 0 | 0 | 0 | 0 | 0 | 0 | 0 | 94,57 | KF036672 | *Dioszegia takashimae* | Basidiomycetous yeast; air sample; contaminated aquifer sediment; sulfide-rich Zodletone spring; plant leaves | NCBI: Related HITs source |
| OTU_773 | 0 | 4 | 0 | 0 | 0 | 0 | 0 | 0 | 0 | 0 | 0 | 91,42 | KC672946 | Uncultured fungus | Isolated from skin, plantar hell. Host = "Homo sapiens". | Findley, K., Oh, J., Yang, J., Conlan, S., Deming, C., Meyer, J. A., … Segre, J. A. (2013). Human Skin Fungal Diversity. Nature, 498(7454), 367–370. http://doi.org/10.1038/nature12171 |
| OTU_8393 | 0 | 0 | 5 | 0 | 0 | 0 | 0 | 0 | 0 | 0 | 0 | 96,40 | KJ708404 | *Sporobolomyces phaffii* | Found in indoor habitats. | Ronald E. Gots , Nancy J. Layton & Suellen W. Pirages (2003): Indoor Health: Background Levels of Fungi, AIHA Journal, 64:4, 427-438. |

La Juanita (LJ); Chascomús (CH); La Limpia (LI); Mar Chiquita (MA2); Vitel (VI); Los Quilmes (QU); - S, corresponded to sorting samples; % Identity is the percent of identity for the OTU taxonomy assignation.

Supplementary Table 5: Pigmented and Non - Pigmented classification for Chrysophyceae, Dicthyophyceae and Dinophyceae classes.

| | PPE Samples | | | | | | FTP Samples | | | | | PR2 Assignation | | | | NCBI - BLAST Revision | | | | Type |
|---|---|---|---|---|---|---|---|---|---|---|---|---|---|---|---|---|---|---|---|---|
| OTU | LJ - S | CH - S | LI - S | MA2 - S | VI - S | QU - S | LJ | CH | LI | MA2 | VI | % of Identity | Phylum | Class | Specie | Accn | % of Identity | Hit description | E-Value | [Pigmented/Non-Pigmented] |
| OTU_11 | 0 | 0 | 0 | 0 | 0 | 4 | 3 | 7 | 0 | 0 | 3159 | 95.26 | Stramenopiles | Chrysophyceae-Synurophyceae | Chrysosphaerella+sp. | HF549059 | 95.27 | Chrysosphaerella brevispina partial 18S rRNA gene, strain S 74.D5 | 0.0 | Pigmented |
| OTU_42 | 0 | 0 | 0 | 0 | 0 | 0 | 1 | 514 | 44 | 14 | 0 | 100 | Stramenopiles | Chrysophyceae-Synurophyceae | Pedospumella+encystans | KX431485 | 100.00 | Pedospumella encystans strain IOW133 18S ribosomal RNA gene, complete sequence | 0.0 | Non-Pigmented |
| OTU_69 | 0 | 0 | 0 | 0 | 0 | 0 | 19 | 194 | 10 | 7 | 49 | 99.05 | Stramenopiles | Chrysophyceae-Synurophyceae | Clade-F_X+sp. | AY651090 | 99.05 | Spumella-like flagellate JBC29 18S ribosomal RNA gene, partial sequence | 0.0 | Non-Pigmented |
| OTU_74 | 0 | 0 | 0 | 0 | 0 | 1 | 363 | 48 | 0 | 5 | 1 | 99.28 | Stramenopiles | Chrysophyceae-Synurophyceae | Clade-C_X+sp. | AY651078 | 99.28 | Spumella-like flagellate JBC22 18S ribosomal RNA gene, partial sequence | 0.0 | Non-Pigmented |
| OTU_92 | 0 | 0 | 0 | 0 | 65 | 1 | 0 | 61 | 0 | 0 | 272 | 92.74 | Stramenopiles | Chrysophyceae-Synurophyceae | Chrysophyceae-Synurophyceae_XXX+sp. | AY651093 | 92.74 | Spumella-like flagellate JBC27 18S ribosomal RNA gene, partial sequence | 1.03e-170 | Non-Pigmented |
| OTU_101 | 0 | 0 | 0 | 0 | 0 | 0 | 0 | 14 | 180 | 0 | 5 | 91.57 | Stramenopiles | Chrysophyceae-Synurophyceae | Chrysophyceae-Synurophyceae_XXX+sp. | AB749080 | 91.55 | Uncultured chrysophyte gene for 18S ribosomal RNA, partial sequence, clone: Bs-St. 5-31 | 6.27e-163 | Unknown |
| OTU_105 | 0 | 0 | 0 | 0 | 0 | 0 | 0 | 8 | 243 | 0 | 9 | 100 | Stramenopiles | Chrysophyceae-Synurophyceae | Synurales_XX+sp. | JQ967306 | 99.54 | Paraphysomonas sp. 25 JMS-2012 18S ribosomal RNA gene, partial sequence | 0.0 | Non-Pigmented |
| OTU_132 | 0 | 0 | 0 | 1 | 0 | 0 | 0 | 40 | 81 | 0 | 1 | 92.31 | Stramenopiles | Chrysophyceae-Synurophyceae | Chrysophyceae-Synurophyceae_XXX+sp. | AB749075 | 92.31 | Uncultured chrysophyte gene for 18S ribosomal RNA, partial sequence, clone: Bs-St. 4-26 | 6.22e-168 | Unknown |
| OTU_137 | 0 | 0 | 0 | 0 | 0 | 0 | 155 | 0 | 0 | 0 | 0 | 99.31 | Stramenopiles | Chrysophyceae-Synurophyceae | Paraphysomonas+foraminifera | JQ967306 | 99.54 | Paraphysomonas sp. 25 JMS-2012 18S ribosomal RNA gene, partial sequence | 0.0 | Non-Pigmented |
| OTU_153 | 0 | 0 | 0 | 0 | 0 | 0 | 0 | 0 | 0 | 598 | 0 | 99.76 | Stramenopiles | Chrysophyceae-Synurophyceae | Chrysophyceae-Synurophyceae_XXX+sp. | JQ967329 | 95.01 | Paraphysomonas sp. 34 JMS-2012 18S ribosomal RNA gene, partial sequence | 0.0 | Non-Pigmented |
| OTU_156 | 0 | 0 | 0 | 0 | 0 | 0 | 11 | 0 | 133 | 4 | 0 | 99.5 | Stramenopiles | Chrysophyceae-Synurophyceae | Clade-C_X+sp. | AB585964 | 93.84 | Spumella sp. TGS1 gene for 18S ribosomal RNA, partial sequence | 2.21e-177 | Non-Pigmented |
| OTU_173 | 0 | 0 | 0 | 0 | 0 | 0 | 0 | 74 | 0 | 13 | 17 | 97.39 | Stramenopiles | Chrysophyceae-Synurophyceae | Clade-F_X+sp. | JQ967323 | 94.79 | Paraphysomonas sp. 18 JMS-2012 18S ribosomal RNA gene, partial sequence | 0.0 | Non-Pigmented |
| OTU_174 | 0 | 0 | 0 | 0 | 0 | 0 | 0 | 43 | 30 | 1 | 0 | 97.14 | Stramenopiles | Chrysophyceae-Synurophyceae | Clade-C_X+sp. | KX431475 | 96.92 | Spumella sp. strain IOW111 18S ribosomal RNA gene, complete sequence | 0.0 | Non-Pigmented |
| OTU_189 | 0 | 0 | 0 | 0 | 0 | 1 | 0 | 0 | 0 | 0 | 169 | 99.52 | Stramenopiles | Chrysophyceae-Synurophyceae | Paraphysomonas+sp. | JQ967330 | 99.53 | Paraphysomonas sp. 22 JMS-2012 18S ribosomal RNA gene, partial sequence | 0.0 | Non-Pigmented |
| OTU_190 | 0 | 0 | 0 | 0 | 0 | 0 | 0 | 20 | 62 | 0 | 0 | 99.29 | Stramenopiles | Chrysophyceae-Synurophyceae | Chrysophyceae-Synurophyceae_XXX+sp. | AB749075 | 99.29 | Uncultured chrysophyte gene for 18S ribosomal RNA, partial sequence, clone: Bs-St. 4-26 | 0.0 | Unknown |
| OTU_194 | 0 | 0 | 0 | 0 | 0 | 0 | 0 | 28 | 7 | 2 | 0 | 99.05 | Stramenopiles | Chrysophyceae-Synurophyceae | Clade-F_X+sp. | AF109326 | 96.91 | Paraphysomonas butcheri 18S small subunit ribosomal RNA gene, complete sequence | 0.0 | Non-Pigmented |
| OTU_196 | 0 | 0 | 0 | 63 | 0 | 0 | 6 | 25 | 47 | 3 | 5 | 98.81 | Stramenopiles | Chrysophyceae-Synurophyceae | Clade-C_X+sp. | KX100620 | 99.05 | Poteriospumella sp. strain D-7-A2 18S ribosomal RNA gene, partial sequence | 0.0 | Non-Pigmented |
| OTU_199 | 0 | 0 | 0 | 0 | 61 | 0 | 6 | 69 | 1 | 4 | 91 | 99.53 | Stramenopiles | Chrysophyceae-Synurophyceae | Clade-F_X+sp. | JQ967313 | 99.53 | Paraphysomonas sp. 33 JMS-2012 18S ribosomal RNA gene, partial sequence | 0.0 | Non-Pigmented |
| OTU_200 | 0 | 0 | 0 | 17 | 0 | 0 | 0 | 2 | 159 | 15 | 0 | 98.02 | Stramenopiles | Chrysophyceae-Synurophyceae | Clade-C_X+sp. | EF165132 | 95.94 | Uroglena sp. CCMP2768 18S ribosomal RNA gene, partial sequence | 0.0 | Pigmented |
| OTU_205 | 0 | 0 | 0 | 1 | 0 | 0 | 0 | 1 | 94 | 0 | 0 | 93.41 | Stramenopiles | Chrysophyceae-Synurophyceae | Chrysophyceae-Synurophyceae_XXX+sp. | AB749080 | 93.41 | Uncultured chrysophyte gene for 18S ribosomal RNA, partial sequence, clone: Bs-St. 5-31 | 2.85e-176 | Unknown |
| OTU_223 | 0 | 0 | 0 | 0 | 0 | 0 | 0 | 39 | 0 | 0 | 4 | 99.76 | Stramenopiles | Chrysophyceae-Synurophyceae | Chrysophyceae-Synurophyceae_XXX+sp. | AB749080 | 99.77 | Uncultured chrysophyte gene for 18S ribosomal RNA, partial sequence, clone: Bs-St. 5-31 | 0.0 | Unknown |
| OTU_232 | 0 | 0 | 0 | 0 | 0 | 0 | 0 | 0 | 0 | 173 | 28 | 97.18 | Stramenopiles | Chrysophyceae-Synurophyceae | Chrysophyceae-Synurophyceae_XXX+sp. | AY651093 | 97.18 | Spumella-like flagellate JBC27 18S ribosomal RNA gene, partial sequence | 0.0 | Non-Pigmented |
| OTU_235 | 0 | 0 | 0 | 0 | 0 | 0 | 71 | 0 | 0 | 2 | 7 | 95.73 | Stramenopiles | Chrysophyceae-Synurophyceae | Clade-F_X+sp. | AY651090 | 95.74 | Spumella-like flagellate JBC29 18S ribosomal RNA gene, partial sequence | 0.0 | Non-Pigmented |
| OTU_237 | 0 | 0 | 0 | 0 | 0 | 0 | 0 | 0 | 0 | 199 | 0 | 99.29 | Stramenopiles | Chrysophyceae-Synurophyceae | Mallomonas+annulata | JQ281516 | 99.53 | Mallomonas annulata strain CCMP474 18S ribosomal RNA gene, partial sequence | 0.0 | Pigmented |
| OTU_238 | 0 | 0 | 0 | 0 | 0 | 0 | 0 | 27 | 41 | 0 | 0 | 97.39 | Stramenopiles | Chrysophyceae-Synurophyceae | Clade-I_X+sp. | KX431464 | 97.39 | Chrysophyceae sp. strain IOW65 18S ribosomal RNA gene, complete sequence | 0.0 | Unknown |
| OTU_249 | 119 | 0 | 0 | 0 | 0 | 0 | 0 | 0 | 0 | 1 | 0 | 98.57 | Stramenopiles | Chrysophyceae-Synurophyceae | Paraphysomonas+sp. | JQ967321 | 98.81 | Paraphysomonas sp. 14 JMS-2012 18S ribosomal RNA gene, partial sequence | 0.0 | Non-Pigmented |
| OTU_273 | 0 | 0 | 0 | 0 | 0 | 0 | 0 | 27 | 6 | 0 | 4 | 92.22 | Stramenopiles | Chrysophyceae-Synurophyceae | Clade-C_X+sp. | EF165111 | 90.85 | Ochromonas vasocystis strain CCMP2741 18S ribosomal RNA gene, partial sequence | 8.18e-157 | Pigmented |
| OTU_286 | 0 | 0 | 0 | 0 | 0 | 0 | 0 | 9 | 0 | 3 | 5 | 97.42 | Stramenopiles | Chrysophyceae-Synurophyceae | Chrysophyceae-Synurophyceae_XXX+sp. | AY651093 | 97.42 | Spumella-like flagellate JBC27 18S ribosomal RNA gene, partial sequence | 0.0 | Non-Pigmented |
| OTU_301 | 0 | 0 | 0 | 0 | 0 | 0 | 0 | 14 | 0 | 0 | 0 | 98.81 | Stramenopiles | Chrysophyceae-Synurophyceae | Clade-C_X+sp. | DQ388542 | 98.81 | Spumella-like flagellate JBNA46 18S ribosomal RNA gene, partial sequence | 0.0 | Non-Pigmented |
| OTU_304 | 0 | 0 | 0 | 0 | 0 | 0 | 0 | 17 | 0 | 0 | 0 | 99.29 | Stramenopiles | Chrysophyceae-Synurophyceae | Clade-H_X+sp. | AB771897 | 99.76 | Uncultured freshwater eukaryote gene for 18S rRNA, partial sequence, clone: K4JAN2012 | 0.0 | Unknown |
| OTU_313 | 0 | 0 | 0 | 0 | 0 | 0 | 5 | 8 | 31 | 1 | 30 | 97.62 | Stramenopiles | Chrysophyceae-Synurophyceae | Clade-F_X+sp. | AY821968 | 97.63 | Uncultured Paraphrysomonas-like chrysophyte clone CV1_B1_34 small subunit ribosomal RNA gene, partial sequence | 0.0 | Non-Pigmented |
| OTU_325 | 0 | 0 | 0 | 0 | 0 | 0 | 7 | 0 | 32 | 0 | 1 | 94.77 | Stramenopiles | Chrysophyceae-Synurophyceae | Chrysophyceae-Synurophyceae_XXX+sp. | JQ967329 | 92.91 | Paraphysomonas sp. 34 JMS-2012 18S ribosomal RNA gene, partial sequence | 1.04e-170 | Non-Pigmented |
| OTU_342 | 0 | 0 | 0 | 0 | 0 | 0 | 0 | 0 | 0 | 88 | 0 | 99.76 | Stramenopiles | Chrysophyceae-Synurophyceae | Clade-F_X+sp. | JQ967319 | 99.76 | Paraphysomonas sp. WA32KAG 18S ribosomal RNA gene, partial sequence | 0.0 | Non-Pigmented |
| OTU_363 | 0 | 0 | 0 | 0 | 0 | 0 | 0 | 47 | 0 | 0 | 0 | 95.49 | Stramenopiles | Chrysophyceae-Synurophyceae | Clade-C_X+sp. | KX431475 | 95.97 | Spumella sp. strain IOW111 18S ribosomal RNA gene, complete sequence | 0.0 | Non-Pigmented |
| OTU_366 | 0 | 0 | 0 | 0 | 0 | 0 | 0 | 17 | 0 | 0 | 0 | 95.98 | Stramenopiles | Chrysophyceae-Synurophyceae | Clade-F_X+sp. | KM516202 | 93.85 | Paraphysomonas sp. SIOpierPara1 18S ribosomal RNA gene, partial sequence | 1.71e-178 | Non-Pigmented |
| OTU_375 | 0 | 0 | 0 | 0 | 0 | 0 | 0 | 4 | 0 | 0 | 0 | 93.6 | Stramenopiles | Chrysophyceae-Synurophyceae | Clade-C_X+sp. | AY082987 | 93.60 | Uncultured eukaryote clone RT5iin35 18S ribosomal RNA gene, complete sequence | 3.70e-175 | Unknown |
| OTU_395 | 0 | 0 | 0 | 0 | 0 | 0 | 0 | 1 | 7 | 9 | 36 | 98.1 | Stramenopiles | Chrysophyceae-Synurophyceae | Clade-F_X+sp. | Z38025 | 94.08 | P.foraminifera ribosomal RNA fragment | 1.32e-179 | Non-Pigmented |
| OTU_414 | 0 | 0 | 0 | 0 | 0 | 0 | 0 | 15 | 0 | 1 | 0 | 98.57 | Stramenopiles | Chrysophyceae-Synurophyceae | Spumella+vulgaris | DQ388563 | 98.58 | Spumella-like flagellate 1031 18S ribosomal RNA gene, partial sequence | 0.0 | Non-Pigmented |
| OTU_430 | 0 | 0 | 0 | 0 | 0 | 0 | 0 | 0 | 13 | 0 | 0 | 94.35 | Stramenopiles | Chrysophyceae-Synurophyceae | Chrysophyceae-Synurophyceae_XXX+sp. | AB749080 | 94.35 | Uncultured chrysophyte gene for 18S ribosomal RNA, partial sequence, clone: Bs-St. 5-31 | 0.0 | Unknown |
| OTU_431 | 0 | 0 | 0 | 0 | 0 | 0 | 0 | 16 | 0 | 0 | 0 | 99.76 | Stramenopiles | Chrysophyceae-Synurophyceae | Chrysophyceae-Synurophyceae_XXX+sp. | KM590581 | 94.10 | Synura sp. JIK-2015b strain Sinsan101709B 18S ribosomal RNA gene, partial sequence | 4.75e-179 | Pigmented |
| OTU_439 | 0 | 0 | 0 | 0 | 0 | 0 | 0 | 0 | 17 | 0 | 0 | 93.33 | Stramenopiles | Chrysophyceae-Synurophyceae | Clade-C_X+sp. | EF165132 | 91.92 | Uroglena sp. CCMP2768 18S ribosomal RNA gene, partial sequence | 4.86e-164 | Pigmented |
| OTU_451 | 0 | 0 | 0 | 0 | 0 | 0 | 0 | 0 | 11 | 0 | 17 | 99.53 | Stramenopiles | Chrysophyceae-Synurophyceae | Synurales_XX+sp. | GU290087 | 99.53 | Uncultured eukaryote clone TKR07M.13 18S ribosomal RNA gene, partial sequence | 0.0 | Unknown |
| OTU_461 | 0 | 0 | 0 | 0 | 0 | 0 | 0 | 2 | 23 | 0 | 1 | 99.3 | Stramenopiles | Chrysophyceae-Synurophyceae | Chrysophyceae-Synurophyceae_XXX+sp. | AY821967 | 99.30 | Uncultured chrysophyte clone CH1_S1_35 small subunit ribosomal RNA gene, partial sequence | 0.0 | Unknown |
| OTU_463 | 0 | 0 | 0 | 0 | 0 | 0 | 0 | 10 | 0 | 13 | 0 | 98.57 | Stramenopiles | Chrysophyceae-Synurophyceae | Clade-F_X+sp. | JQ967325 | 98.81 | Paraphysomonas sp. 20 JMS-2012 18S ribosomal RNA gene, partial sequence | 0.0 | Non-Pigmented |
| OTU_497 | 0 | 0 | 0 | 0 | 0 | 0 | 0 | 4 | 0 | 0 | 0 | 97.18 | Stramenopiles | Chrysophyceae-Synurophyceae | Chrysophyceae-Synurophyceae_XXX+sp. | AB749075 | 97.18 | Uncultured chrysophyte gene for 18S ribosomal RNA, partial sequence, clone: Bs-St. 4-26 | 0.0 | Unknown |
| OTU_516 | 0 | 0 | 0 | 0 | 0 | 0 | 0 | 6 | 0 | 0 | 0 | 97.17 | Stramenopiles | Chrysophyceae-Synurophyceae | Synurales_XX+sp. | AB622331 | 97.17 | Uncultured freshwater eukaryote gene for 18S rRNA, partial sequence, clone: K6APR2010 | 0.0 | Unknown |
| OTU_518 | 0 | 0 | 0 | 0 | 0 | 0 | 0 | 3 | 0 | 0 | 0 | 95.27 | Stramenopiles | Chrysophyceae-Synurophyceae | Chrysophyceae-Synurophyceae_XXX+sp. | KM817864 | 92.89 | Mallomonas ceylanica strain Gungnam092213J 18S ribosomal RNA gene, partial sequence | 2.88e-171 | Pigmented |
| OTU_525 | 0 | 0 | 0 | 0 | 0 | 0 | 0 | 5 | 0 | 0 | 0 | 98.33 | Stramenopiles | Chrysophyceae-Synurophyceae | Clade-C_X+sp. | EF165132 | 95.95 | Uroglena sp. CCMP2768 18S ribosomal RNA gene, partial sequence | 0.0 | Pigmented |
| OTU_551 | 0 | 0 | 0 | 0 | 0 | 0 | 10 | 0 | 0 | 2 | 4 | 99.29 | Stramenopiles | Chrysophyceae-Synurophyceae | Paraphysomonas+butcheri | JQ967291 | 99.29 | Paraphysomonas butcheri isolate MD03 18S ribosomal RNA gene, partial sequence | 0.0 | Non-Pigmented |
| OTU_587 | 0 | 0 | 0 | 0 | 0 | 0 | 0 | 0 | 0 | 9 | 0 | 99.05 | Stramenopiles | Chrysophyceae-Synurophyceae | Mallomonas+caudata | GU935629 | 99.52 | Mallomonas caudata strain Dangje060207A 18S ribosomal RNA gene, partial sequence | 0.0 | Pigmented |
| OTU_590 | 0 | 0 | 0 | 0 | 0 | 0 | 0 | 0 | 0 | 12 | 0 | 98.57 | Stramenopiles | Chrysophyceae-Synurophyceae | Uroglena+sp. | EF165132 | 98.57 | Uroglena sp. CCMP2768 18S ribosomal RNA gene, partial sequence | 0.0 | Pigmented |
| OTU_643 | 0 | 0 | 0 | 0 | 0 | 0 | 0 | 0 | 0 | 0 | 4 | 100 | Stramenopiles | Chrysophyceae-Synurophyceae | Clade-C_X+sp. | EF165132 | 96.18 | Uroglena sp. CCMP2768 18S ribosomal RNA gene, partial sequence | 0.0 | Pigmented |
| OTU_685 | 0 | 0 | 0 | 0 | 0 | 0 | 0 | 3 | 0 | 2 | 0 | 96.67 | Stramenopiles | Chrysophyceae-Synurophyceae | Chrysophyceae-Synurophyceae_XXX+sp. | JQ967331 | 94.54 | Paraphysomonas sp. 21 JMS-2012 18S ribosomal RNA gene, partial sequence | 0.0 | Non-Pigmented |
| OTU_693 | 0 | 0 | 0 | 0 | 0 | 0 | 0 | 0 | 0 | 0 | 0 | 96.21 | Stramenopiles | Chrysophyceae-Synurophyceae | Chrysosphaerella+sp. | JQ281516 | 93.62 | Mallomonas annulata strain CCMP474 18S ribosomal RNA gene, partial sequence | 2.85e-176 | Pigmented |
| OTU_727 | 0 | 0 | 0 | 0 | 0 | 0 | 0 | 2 | 0 | 0 | 2 | 91.53 | Stramenopiles | Chrysophyceae-Synurophyceae | Clade-F_X+sp. | GU822961 | 91.57 | Uncultured stramenopile clone AB3F14RJ3E10 18S ribosomal RNA gene, partial sequence | 2.25e-162 | Unknown |
| OTU_746 | 0 | 0 | 0 | 0 | 0 | 0 | 0 | 2 | 0 | 0 | 0 | 94.84 | Stramenopiles | Chrysophyceae-Synurophyceae | Chrysophyceae-Synurophyceae_XXX+sp. | AB749075 | 94.84 | Uncultured chrysophyte gene for 18S ribosomal RNA, partial sequence, clone: Bs-St. 4-26 | 0.0 | Unknown |
| OTU_770 | 0 | 0 | 0 | 0 | 0 | 0 | 0 | 3 | 1 | 0 | 0 | 97.86 | Stramenopiles | Chrysophyceae-Synurophyceae | Clade-F_X+sp. | JQ967323 | 92.89 | Paraphysomonas sp. 18 JMS-2012 18S ribosomal RNA gene, partial sequence | 1.03e-170 | Non-Pigmented |
| OTU_791 | 0 | 0 | 0 | 0 | 0 | 0 | 0 | 0 | 0 | 4 | 0 | 99.76 | Stramenopiles | Chrysophyceae-Synurophyceae | Synura+petersenii | KY464904 | 99.76 | Synura sp. strain WA18KA 18S ribosomal RNA gene, partial sequence | 0.0 | Pigmented |
| OTU_797 | 0 | 0 | 0 | 0 | 0 | 0 | 0 | 0 | 0 | 0 | 6 | 95.72 | Stramenopiles | Chrysophyceae-Synurophyceae | Paraphysomonas+sp. | JQ967330 | 95.72 | Paraphysomonas sp. 22 JMS-2012 18S ribosomal RNA gene, partial sequence | 0.0 | Non-Pigmented |
| OTU_862 | 0 | 0 | 0 | 0 | 0 | 0 | 0 | 19 | 0 | 0 | 0 | 96.42 | Stramenopiles | Chrysophyceae-Synurophyceae | Uroglena+sp. | EF165132 | 96.42 | Uroglena sp. CCMP2768 18S ribosomal RNA gene, partial sequence | 0.0 | Pigmented |
| OTU_925 | 0 | 0 | 0 | 0 | 0 | 0 | 0 | 0 | 0 | 0 | 0 | 99.05 | Stramenopiles | Chrysophyceae-Synurophyceae | Oikomonas+sp. | AY520451 | 99.05 | Oikomonas sp. SA-2.2 18S ribosomal RNA gene, partial sequence | 0.0 | Non-Pigmented |
| OTU_934 | 0 | 0 | 0 | 0 | 0 | 0 | 0 | 1 | 8 | 0 | 0 | 94.27 | Stramenopiles | Chrysophyceae-Synurophyceae | Uroglena+sp. | EF165132 | 94.27 | Uroglena sp. CCMP2768 18S ribosomal RNA gene, partial sequence | 3.67e-180 | Pigmented |
| OTU_985 | 0 | 0 | 0 | 0 | 0 | 0 | 1 | 25 | 0 | 0 | 0 | 98.1 | Stramenopiles | Chrysophyceae-Synurophyceae | Clade-C_X+sp. | AJ236861 | 97.86 | Spumella danica 18S rDNA gene | 0.0 | Non-Pigmented |
| OTU_1011 | 0 | 0 | 0 | 0 | 1 | 0 | 0 | 0 | 0 | 0 | 0 | 97.88 | Stramenopiles | Chrysophyceae-Synurophyceae | Clade-F_X+sp. | JQ967313 | 97.88 | Paraphysomonas sp. 33 JMS-2012 18S ribosomal RNA gene, partial sequence | 4.05e-110 | Non-Pigmented |
| OTU_1013 | 0 | 0 | 0 | 0 | 0 | 0 | 0 | 1 | 0 | 0 | 0 | 97.15 | Stramenopiles | Chrysophyceae-Synurophyceae | Clade-C_X+sp. | KX100620 | 96.44 | Poteriospumella sp. strain D-7-A2 18S ribosomal RNA gene, partial sequence | 0.0 | Non-Pigmented |
| OTU_1018 | 0 | 0 | 0 | 0 | 0 | 0 | 0 | 0 | 0 | 0 | 0 | 87.03 | Stramenopiles | Chrysophyceae-Synurophyceae | Clade-F_X+sp. | Unknown | Unknown | Unknown | Unknown | Unknown |
| OTU_1040 | 0 | 0 | 0 | 0 | 0 | 0 | 0 | 0 | 0 | 11 | 0 | 98.1 | Stramenopiles | Chrysophyceae-Synurophyceae | Chrysolepidomonas+dendrolepidota | AF123297 | 98.10 | Chrysolepidomonas dendrolepidota small subunit ribosomal RNA gene, partial sequence | 0.0 | Pigmented |
| OTU_1088 | 0 | 0 | 0 | 0 | 0 | 0 | 0 | 5 | 0 | 4 | 17 | 96.25 | Stramenopiles | Chrysophyceae-Synurophyceae | Chrysophyceae-Synurophyceae_XXX+sp. | AY651093 | 96.25 | Spumella-like flagellate JBC27 18S ribosomal RNA gene, partial sequence | 0.0 | Non-Pigmented |
| OTU_1128 | 0 | 0 | 0 | 0 | 0 | 0 | 0 | 5 | 0 | 0 | 0 | 97.15 | Stramenopiles | Chrysophyceae-Synurophyceae | Clade-F_X+sp. | AF109326 | 95.72 | Paraphysomonas butcheri 18S small subunit ribosomal RNA gene, complete sequence | 0.0 | Non-Pigmented |
| OTU_1139 | 0 | 0 | 0 | 0 | 0 | 0 | 0 | 1 | 0 | 0 | 0 | 97.09 | Stramenopiles | Chrysophyceae-Synurophyceae | Synurales_XX+sp. | GU290087 | 96.45 | Uncultured eukaryote clone TKR07M.13 18S ribosomal RNA gene, partial sequence | 0.0 | Unknown |
| OTU_1174 | 0 | 0 | 0 | 0 | 0 | 0 | 0 | 0 | 0 | 7 | 0 | 98.8 | Stramenopiles | Chrysophyceae-Synurophyceae | Clade-C_X+sp. | AY651077 | 99.04 | Spumella-like flagellate JBAF33 18S ribosomal RNA gene, partial sequence | 0.0 | Non-Pigmented |
| OTU_1265 | 0 | 0 | 0 | 0 | 0 | 0 | 0 | 0 | 0 | 15 | 0 | 94.12 | Stramenopiles | Chrysophyceae-Synurophyceae | Chrysophyceae-Synurophyceae_XXX+sp. | LN581250 | 92.02 | Uncultured eukaryote partial 18S rRNA gene, clone SIBO712_N12D1_18S_E | 6.97e-88 | Unknown |
| OTU_1287 | 0 | 0 | 0 | 0 | 0 | 0 | 0 | 1 | 0 | 0 | 0 | 94.13 | Stramenopiles | Chrysophyceae-Synurophyceae | Chrysophyceae-Synurophyceae_XXX+sp. | AB749080 | 94.13 | Uncultured chrysophyte gene for 18S ribosomal RNA, partial sequence, clone: Bs-St. 5-31 | 0.0 | Unknown |
| OTU_1339 | 0 | 0 | 0 | 0 | 0 | 0 | 0 | 0 | 0 | 5 | 0 | 96.45 | Stramenopiles | Chrysophyceae-Synurophyceae | Chrysophyceae-Synurophyceae_XXX+sp. | KX100597 | 92.64 | Paraphysomonas sp. strain PR26KA 18S ribosomal RNA gene, partial sequence | 1.34e-169 | Non-Pigmented |
| OTU_1387 | 0 | 0 | 0 | 0 | 0 | 0 | 0 | 0 | 0 | 0 | 0 | 93.16 | Stramenopiles | Chrysophyceae-Synurophyceae | Clade-C_X+sp. | EF165132 | 91.29 | Uroglena sp. CCMP2768 18S ribosomal RNA gene, partial sequence | 1.36e-159 | Pigmented |
| OTU_1469 | 0 | 0 | 0 | 0 | 0 | 0 | 0 | 0 | 0 | 0 | 3 | 96.41 | Stramenopiles | Chrysophyceae-Synurophyceae | Clade-C_X+sp. | KX431505 | 93.35 | Spumella sp. strain IOW85 18S ribosomal RNA gene, partial sequence | 1.73e-173 | Non-Pigmented |
| OTU_1493 | 0 | 0 | 0 | 0 | 0 | 0 | 0 | 10 | 0 | 0 | 0 | 96.96 | Stramenopiles | Chrysophyceae-Synurophyceae | Chrysophyceae-Synurophyceae_XXX+sp. | AY651093 | 96.96 | Spumella-like flagellate JBC27 18S ribosomal RNA gene, partial sequence | 0.0 | Non-Pigmented |
| OTU_1498 | 0 | 0 | 0 | 0 | 0 | 0 | 0 | 0 | 0 | 1 | 18 | 96.67 | Stramenopiles | Chrysophyceae-Synurophyceae | Paraphysomonas+butcheri | AF109326 | 97.15 | Paraphysomonas butcheri 18S small subunit ribosomal RNA gene, complete sequence | 0.0 | Non-Pigmented |
| OTU_1527 | 0 | 0 | 0 | 0 | 0 | 0 | 0 | 0 | 0 | 0 | 0 | 82 | Stramenopiles | Chrysophyceae-Synurophyceae | Leukarachnion+sp. | Unknown | Unknown | Unknown | Unknown | Unknown |
| OTU_3154 | 0 | 0 | 0 | 0 | 0 | 0 | 0 | 0 | 12 | 0 | 4 | 97.39 | Stramenopiles | Chrysophyceae-Synurophyceae | Clade-F_X+sp. | AF174376 | 93.37 | Paraphysomonas foraminifera 18S ribosomal RNA gene, partial sequence | 1.33e-174 | Non-Pigmented |
| OTU_4398 | 0 | 0 | 0 | 2 | 0 | 0 | 0 | 0 | 4 | 0 | 0 | 95.3 | Stramenopiles | Chrysophyceae-Synurophyceae | Clade-C_X+sp. | DQ388542 | 93.81 | Spumella-like flagellate JBNA46 18S ribosomal RNA gene, partial sequence | 7.98e-177 | Non-Pigmented |
| OTU_5059 | 0 | 0 | 0 | 0 | 0 | 0 | 0 | 0 | 0 | 0 | 3 | 97.39 | Stramenopiles | Chrysophyceae-Synurophyceae | Clade-F_X+sp. | Z38025 | 93.84 | P.foraminifera ribosomal RNA fragment | 6.17e-178 | Non-Pigmented |
| OTU_9494 | 0 | 0 | 0 | 0 | 0 | 0 | 0 | 3 | 0 | 1 | 0 | 97.62 | Stramenopiles | Chrysophyceae-Synurophyceae | Clade-F_X+sp. | Z38025 | 93.62 | P.foraminifera ribosomal RNA fragment | 2.87e-176 | Non-Pigmented |
| OTU_9581 | 0 | 0 | 0 | 0 | 0 | 0 | 0 | 0 | 0 | 0 | 12 | 96.67 | Stramenopiles | Chrysophyceae-Synurophyceae | Paraphysomonas+butcheri | AF109326 | 97.15 | Paraphysomonas butcheri 18S small subunit ribosomal RNA gene, complete sequence | 0.0 | Non-Pigmented |
| OTU_9878 | 0 | 0 | 0 | 0 | 0 | 0 | 0 | 4 | 0 | 6 | 0 | 97.62 | Stramenopiles | Chrysophyceae-Synurophyceae | Clade-F_X+sp. | Z38025 | 94.09 | P.foraminifera ribosomal RNA fragment | 1.33e-179 | Non-Pigmented |
| OTU_11233 | 0 | 0 | 0 | 0 | 0 | 0 | 0 | 0 | 0 | 0 | 0 | 95.01 | Stramenopiles | Chrysophyceae-Synurophyceae | Clade-F_X+sp. | JQ967323 | 92.67 | Paraphysomonas sp. 18 JMS-2012 18S ribosomal RNA gene, partial sequence | 1.34e-169 | Non-Pigmented |
| OTU_14190 | 0 | 0 | 0 | 0 | 0 | 0 | 0 | 2 | 0 | 0 | 5 | 91.8 | Stramenopiles | Chrysophyceae-Synurophyceae | Chrysophyceae-Synurophyceae_XXX+sp. | AB749075 | 91.80 | Uncultured chrysophyte gene for 18S ribosomal RNA, partial sequence, clone: Bs-St. 4-26 | 1.35e-164 | Unknown |
| OTU_253 | 0 | 0 | 0 | 0 | 0 | 0 | 27 | 27 | 0 | 2 | 0 | 98.83 | Stramenopiles | Dictyochophyceae | Pedinellales_X+sp. | JF794054 | 96.04 | Pedinellales sp. RCC2289 18S ribosomal RNA gene, partial sequence | 0.0 | Pigmented |
| OTU_309 | 0 | 0 | 0 | 0 | 0 | 0 | 0 | 19 | 22 | 0 | 3 | 96.8 | Stramenopiles | Dictyochophyceae | Pedinellales_X+sp. | JF794054 | 90.66 | Pedinellales sp. RCC2289 18S ribosomal RNA gene, partial sequence | 4.90e-159 | Pigmented |
| OTU_487 | 0 | 0 | 0 | 0 | 0 | 0 | 0 | 1 | 0 | 0 | 6 | 96.13 | Stramenopiles | Dictyochophyceae | Pedinellales_X+sp. | EU024988 | 91.59 | Pseudopedinella sp. FU44-42M 18S ribosomal RNA gene, partial sequence | 8.09e-167 | Pigmented |
| OTU_825 | 0 | 0 | 0 | 0 | 0 | 0 | 1 | 2 | 0 | 1 | 0 | 99.77 | Stramenopiles | Dictyochophyceae | Pedinellales_X+sp. | JF794054 | 96.50 | Pedinellales sp. RCC2289 18S ribosomal RNA gene, partial sequence | 0.0 | Pigmented |
| OTU_983 | 0 | 0 | 0 | 0 | 0 | 0 | 0 | 0 | 0 | 0 | 0 | 87.24 | Stramenopiles | Dictyochophyceae | Pedinellales_X+sp. | AM901361 | 97.07 | Cryptomonas sp. M1634 partial 18S rRNA gene, strain M1634 | 6.81e-108 | Pigmented |
| OTU_44 | 0 | 0 | 0 | 0 | 0 | 2 | 0 | 0 | 0 | 2 | 1873 | 84.2 | Alveolata | Dinophyceae | Suessiales_XX+sp. | FJ903098 | 98.39 | Uncultured dinoflagellate clone LP80ME80 18S ribosomal RNA gene, partial sequence | 0.0 | Unknown |
| OTU_57 | 0 | 0 | 0 | 0 | 0 | 0 | 0 | 357 | 5 | 12 | 0 | 97.37 | Alveolata | Dinophyceae | Prorocentrum+sp. | KX465200 | 97.85 | Uncultured dinoflagellate clone OL56 18S ribosomal RNA gene, partial sequence | 0.0 | Unknown |
| OTU_142 | 14 | 0 | 0 | 0 | 0 | 0 | 0 | 233 | 0 | 3 | 22 | 99.28 | Alveolata | Dinophyceae | Pfiesteriaceae_X+sp. | KR535602 | 98.09 | Scrippsiella trochoidea clone stain BDH 18S ribosomal RNA gene, partial sequence | 0.0 | Pigmented |
| OTU_211 | 0 | 0 | 0 | 0 | 0 | 0 | 0 | 0 | 96 | 0 | 5 | 96.41 | Alveolata | Dinophyceae | Woloszynskia+leopoliensis | JQ639766 | 96.66 | Woloszynskia sp. QZ-2012 voucher HBI:SC201101a small subunit ribosomal RNA gene, partial sequence | 0.0 | Pigmented |
| OTU_254 | 0 | 0 | 0 | 0 | 0 | 0 | 0 | 0 | 49 | 0 | 0 | 95.95 | Alveolata | Dinophyceae | Prorocentrum+sp. | HQ191345 | 95.95 | Uncultured eukaryote clone PA2009C9 18S ribosomal RNA gene, internal transcribed spacer 1, 5.8S ribosomal RNA gene, and internal transcribed spacer 2, complete sequence; and 28S ribosomal RNA gene, partial sequence | 0.0 | Unknown |
| OTU_352 | 0 | 0 | 0 | 0 | 1 | 0 | 0 | 20 | 0 | 1 | 0 | 93.56 | Alveolata | Dinophyceae | Dinophyceae_XXX+sp. | HQ270473 | 93.08 | Gymnodinium sp. NVA/RUS/2008 clone 44 18S ribosomal RNA gene and internal transcribed spacer 1, partial sequence | 7.93e-172 | Unknown |
| OTU_405 | 0 | 0 | 0 | 0 | 0 | 0 | 37 | 0 | 0 | 0 | 0 | 93.06 | Alveolata | Dinophyceae | Prorocentrum+sp. | KJ925246 | 93.06 | Uncultured eukaryote clone 270E12 small subunit ribosomal RNA gene, partial sequence | 1.03e-170 | Unknown |
| OTU_406 | 0 | 0 | 0 | 0 | 0 | 0 | 0 | 0 | 13 | 0 | 0 | 92.84 | Alveolata | Dinophyceae | Prorocentrum+sp. | KJ925246 | 92.84 | Uncultured eukaryote clone 270E12 small subunit ribosomal RNA gene, partial sequence | 1.33e-169 | Unknown |
| OTU_418 | 0 | 0 | 0 | 0 | 0 | 0 | 0 | 0 | 0 | 7 | 25 | 99.28 | Alveolata | Dinophyceae | Scrippsiella+hangoei | HG005132 | 99.28 | Stoeckeria sp. SSSC09 genomic DNA containing 18S rRNA gene, ITS1, 5.8S rRNA gene, ITS2 and 28S rRNA gene, strain SSSC09 | 0.0 | Pigmented |
| OTU_459 | 0 | 0 | 0 | 0 | 0 | 0 | 0 | 2 | 0 | 0 | 0 | 87.12 | Alveolata | Dinophyceae | Dinophyceae_XXX+sp. | Unknown | Unknown | Unknown | Unknown | Unknown |
| OTU_602 | 0 | 0 | 0 | 0 | 0 | 0 | 0 | 0 | 2 | 0 | 8 | 96.66 | Alveolata | Dinophyceae | Prorocentrum+sp. | EF196722 | 99.05 | Uncultured eukaryotic picoplankton clone B26 18S ribosomal RNA gene, partial sequence | 0.0 | Unknown |
| OTU_607 | 0 | 0 | 0 | 0 | 0 | 0 | 0 | 6 | 0 | 0 | 0 | 93.79 | Alveolata | Dinophyceae | Prorocentrum+sp. | JN090897 | 93.80 | Uncultured eukaryote clone KRL01E37 18S ribosomal RNA gene, partial sequence | 1.02e-175 | Unknown |
| OTU_617 | 0 | 0 | 0 | 0 | 0 | 0 | 0 | 86 | 0 | 1 | 115 | 97.85 | Alveolata | Dinophyceae | Prorocentrum+sp. | KJ925246 | 97.85 | Uncultured eukaryote clone 270E12 small subunit ribosomal RNA gene, partial sequence | 0.0 | Unknown |
| OTU_620 | 0 | 0 | 0 | 0 | 0 | 0 | 0 | 0 | 0 | 17 | 0 | 97.13 | Alveolata | Dinophyceae | Prorocentrum+sp. | JN090897 | 97.13 | Uncultured eukaryote clone KRL01E37 18S ribosomal RNA gene, partial sequence | 0.0 | Unknown |
| OTU_629 | 0 | 0 | 0 | 0 | 0 | 0 | 0 | 2 | 0 | 0 | 0 | 99.28 | Alveolata | Dinophyceae | Dinophyceae_XXX+sp. | FJ473378 | 98.33 | Dissodinium pseudolunula isolate FG59 18S ribosomal RNA gene, partial sequence | 0.0 | Non-Pigmented |
| OTU_731 | 0 | 0 | 0 | 0 | 0 | 0 | 0 | 0 | 0 | 24 | 0 | 99.05 | Alveolata | Dinophyceae | Suessiales_XX+sp. | LC068838 | 98.81 | Suessiaceae sp. od12-21kt genes for 18S rRNA, ITS1, 5.8S rRNA, ITS2, 28S rRNA, partial and complete sequence, strain: od12-21kt | 0.0 | Unknown |
| OTU_793 | 0 | 0 | 0 | 0 | 0 | 0 | 0 | 0 | 203 | 73 | 0 | 97.37 | Alveolata | Dinophyceae | Prorocentrum+sp. | JN090873 | 97.37 | Uncultured eukaryote clone KRL01E13 18S ribosomal RNA gene, partial sequence | 0.0 | Unknown |
| OTU_854 | 0 | 0 | 0 | 0 | 0 | 0 | 0 | 0 | 0 | 0 | 12 | 97.37 | Alveolata | Dinophyceae | Prorocentrum+sp. | JN090897 | 97.37 | Uncultured eukaryote clone KRL01E37 18S ribosomal RNA gene, partial sequence | 0.0 | Unknown |
| OTU_904 | 4 | 0 | 0 | 0 | 0 | 0 | 1 | 0 | 0 | 0 | 0 | 85.55 | Alveolata | Dinophyceae | Dinophyceae_XXX+sp. | KX602083 | 94.51 | Uncultured Amoebophrya clone 18_f3_1 small subunit ribosomal RNA gene, partial sequence | 0.0 | Non-Pigmented |
| OTU_982 | 0 | 0 | 0 | 0 | 0 | 0 | 0 | 0 | 0 | 0 | 0 | 90.59 | Alveolata | Dinophyceae | Prorocentrum+sp. | EF196722 | 91.06 | Uncultured eukaryotic picoplankton clone B26 18S ribosomal RNA gene, partial sequence | 1.74e-158 | Unknown |
| OTU_1005 | 0 | 0 | 0 | 0 | 0 | 0 | 0 | 2 | 0 | 0 | 0 | 90.34 | Alveolata | Dinophyceae | Ciliophrys+infusionum | AB846664 | 90.34 | Ciliophrys infusionum DNA, ubiquitin specific peptidase 39, ubiquitin conjugating enzyme 12, pseudogenes of 18S ribosomal RNA, 5.8S ribosomal RNA, 28S ribosomal RNA, strain: YPF1207 | 5.42e-79 | Non-Pigmented |
| OTU_1328 | 0 | 0 | 0 | 0 | 0 | 0 | 0 | 0 | 0 | 5 | 0 | 89.76 | Alveolata | Dinophyceae | Amphidinium+sp. | LC054921 | 95.25 | Amphidinium steinii gene for 18S ribosomal RNA, partial sequence, strain: HG220 | 0.0 | Pigmented |
| OTU_1506 | 0 | 0 | 0 | 0 | 0 | 0 | 0 | 0 | 0 | 0 | 3 | 95 | Alveolata | Dinophyceae | Prorocentrum+sp. | KJ925246 | 95.00 | Uncultured eukaryote clone 270E12 small subunit ribosomal RNA gene, partial sequence | 0.0 | Unknown |
| OTU_1666 | 0 | 0 | 0 | 0 | 0 | 0 | 0 | 0 | 0 | 10 | 0 | 97.85 | Alveolata | Dinophyceae | Suessiales_XX+sp. | LC068843 | 94.51 | Biecheleriopsis adriatica genes for 18S rRNA, ITS1, 5.8S rRNA, ITS2, 28S rRNA, partial and complete sequence, strain: trd278-kt | 0.0 | Pigmented |
| OTU_1727 | 0 | 0 | 0 | 0 | 0 | 0 | 0 | 1 | 0 | 0 | 4 | 96.41 | Alveolata | Dinophyceae | Prorocentrum+sp. | KX465200 | 97.13 | Uncultured dinoflagellate clone OL56 18S ribosomal RNA gene, partial sequence | 0.0 | Unknown |
| OTU_2456 | 0 | 0 | 0 | 0 | 0 | 0 | 0 | 0 | 0 | 6 | 0 | 98.09 | Alveolata | Dinophyceae | Prorocentrum+sp. | KJ925246 | 98.09 | Uncultured eukaryote clone 270E12 small subunit ribosomal RNA gene, partial sequence | 0.0 | Unknown |
| OTU_2814 | 0 | 0 | 0 | 0 | 0 | 0 | 1 | 1 | 0 | 0 | 2 | 96.42 | Alveolata | Dinophyceae | Prorocentrum+sp. | KJ925299 | 96.42 | Uncultured eukaryote clone 272F01 small subunit ribosomal RNA gene, partial sequence | 0.0 | Unknown |
| OTU_3558 | 0 | 0 | 0 | 0 | 0 | 0 | 0 | 0 | 1 | 0 | 2 | 97.62 | Alveolata | Dinophyceae | Pfiesteriaceae_X+sp. | LK934662 | 97.62 | Aduncusdinium glandulum 18S rRNA gene, 5.8S rRNA gene, 28S rRNA gene, ITS1 and ITS2 | 0.0 | Unknown |
| OTU_3835 | 0 | 0 | 0 | 0 | 0 | 0 | 0 | 0 | 2 | 2 | 0 | 96.66 | Alveolata | Dinophyceae | Dinophyceae_XXX+sp. | EF058241 | 99.04 | Peridiniopsis borgei strain PBSK A small subunit ribosomal RNA gene, partial sequence | 0.0 | Pigmented |
| OTU_4258 | 0 | 0 | 0 | 0 | 0 | 0 | 0 | 0 | 0 | 12 | 0 | 94.75 | Alveolata | Dinophyceae | Prorocentrum+sp. | HQ191345 | 94.75 | Uncultured eukaryote clone PA2009C9 18S ribosomal RNA gene, internal transcribed spacer 1, 5.8S ribosomal RNA gene, and internal transcribed spacer 2, complete sequence; and 28S ribosomal RNA gene, partial sequence | 0.0 | Unknown |
| OTU_5432 | 0 | 0 | 0 | 0 | 0 | 0 | 0 | 12 | 0 | 8 | 0 | 96.17 | Alveolata | Dinophyceae | Prorocentrum+sp. | KX465200 | 97.85 | Uncultured dinoflagellate clone OL56 18S ribosomal RNA gene, partial sequence | 0.0 | Unknown |
| OTU_6129 | 0 | 0 | 0 | 0 | 0 | 0 | 0 | 0 | 0 | 0 | 0 | 96.17 | Alveolata | Dinophyceae | Prorocentrum+sp. | KJ925246 | 96.17 | Uncultured eukaryote clone 270E12 small subunit ribosomal RNA gene, partial sequence | 0.0 | Unknown |
| OTU_6649 | 0 | 0 | 0 | 0 | 0 | 0 | 0 | 0 | 0 | 0 | 0 | 96.17 | Alveolata | Dinophyceae | Prorocentrum+sp. | KJ925246 | 96.17 | Uncultured eukaryote clone 270E12 small subunit ribosomal RNA gene, partial sequence | 0.0 | Unknown |
| OTU_6981 | 0 | 0 | 0 | 0 | 0 | 0 | 0 | 1 | 0 | 4 | 1 | 96.17 | Alveolata | Dinophyceae | Prorocentrum+sp. | KX465200 | 97.61 | Uncultured dinoflagellate clone OL56 18S ribosomal RNA gene, partial sequence | 0.0 | Unknown |
| OTU_7068 | 0 | 0 | 0 | 0 | 0 | 0 | 0 | 5 | 0 | 0 | 0 | 95.69 | Alveolata | Dinophyceae | Prorocentrum+sp. | KX465200 | 97.37 | Uncultured dinoflagellate clone OL56 18S ribosomal RNA gene, partial sequence | 0.0 | Unknown |
| OTU_13856 | 0 | 0 | 0 | 0 | 0 | 0 | 0 | 0 | 0 | 0 | 1 | 98.23 | Alveolata | Dinophyceae | Dinophyceae_XXXX | FJ903103 | 98.24 | Uncultured dinoflagellate clone LP80ME90 18S ribosomal RNA gene, partial sequence | 2.87e-166 | Unknown |

La Juanita (LJ); Chascomús (CH); La Limpia (LI); Mar Chiquita (MA2); Vitel (VI); Los Quilmes (QU);

Supplementary Table 6: Number (N°) of reads and OTUs obtained for each sample before samples standardization. The number of shared OTUs corresponds to the number of OTUs that were presents in both cell sorting and FTP samples for each sites of study after standarization.

| Name | ID | N° of reads environmental (FTP) | N° of reads sorting (PPE) | N° of OTUs environmental (FTP) | N° of OTUs sorting (PPE) | N° of shared OTUs |
|---|---|---|---|---|---|---|
| La Juanita | LJ | 157 562 | 90 609 | 204 | 83 | 28 |
| Chascomús | CH | 275 343 | 272 264 | 442 | 84 | 23 |
| La Limpia | LI | 133 243 | 24 402 | 176 | 26 | 11 |
| Mar Chiquita | MA2 | 23 637 | 131 022 | 349 | 64 | 26 |
| Vitel | VI | 78 627 | 66 019 | 398 | 50 | 33 |
| Los Quilmes | QU | - | 40 902 | - | 65 | - |
| Average | | 133 682 | 104 203 | 314 | 62 | 24 |

Supplementary Table 7: Abundant OTUs (i.e. that contribute to more than the 1% of the total number of reads), from, respectively, FTP samples and from cell sorting samples (in red).
Taxonomic assignations for each OTU were shown with their percentage of identity with PR2 reference database.

| | Sorting Samples (PPE) | | | | | | Environmental Samples (FTP) | | | | | | | | | | | |
|---|---|---|---|---|---|---|---|---|---|---|---|---|---|---|---|---|---|---|
| **OTU** | **LJ-S** | **CH-S** | **LI-S** | **MA2-S** | **VI-S** | **QU-S** | **LJ** | **CH** | **LI** | **MA2** | **VI** | **% of Identity** | **Domain** | **Phylum** | **Division** | **Class** | **Genus** | **Specie** |
| OTU_1 | 2670 | 16857 | 734 | 17363 | 10025 | 1914 | 2 | 18 | 17 | 37 | 2 | 99,52 | Eukaryota | Archaeplastida | Chlorophyta | Trebouxiophyceae | Choricystis | Choricystis+minor |
| OTU_2 | 313 | 0 | 1 | 417 | 297 | 6 | 4119 | 413 | 8959 | 5211 | 1925 | 100,00 | Eukaryota | Cryptista | Cryptophyta | Cryptophyceae | Cryptomonas | Cryptomonas+marssonii |
| OTU_3 | 3248 | 1 | 3 | 1 | 11 | 1 | 5247 | 416 | 2627 | 4741 | 510 | 100,00 | Eukaryota | Cryptista | Cryptophyta | Cryptophyceae | Cryptomonas | Cryptomonas+sp. |
| OTU_4 | 62 | 1840 | 2 | 84 | 881 | 20956 | 0 | 38 | 0 | 26 | 10 | 99,28 | Eukaryota | Archaeplastida | Chlorophyta | Chlorophyceae | Mychonastes | Mychonastes+homosphaera |
| OTU_5 | 0 | 15 | 0 | 0 | 0 | 0 | 2 | 2060 | 18 | 240 | 103 | 97,82 | Eukaryota | Alveolata | Perkinsea | Perkinsida | Perkinsida_XXX | Perkinsida_XXX+sp. |
| OTU_7 | 0 | 0 | 0 | 0 | 0 | 0 | 158 | 1891 | 0 | 89 | 64 | 98,57 | Eukaryota | Stramenopiles | Ochrophyta | Bacillariophyta | Cyclotella | Cyclotella+choctawhatcheeana |
| OTU_8 | 0 | 0 | 0 | 0 | 0 | 0 | 1 | 1488 | 27 | 0 | 1 | 91,02 | Eukaryota | Opisthokonta | Choanoflagellida | Choanoflagellatea | Monosigidae_Group_O_X | Monosigidae_Group_O_X+sp. |
| OTU_10 | 0 | 0 | 0 | 0 | 0 | 0 | 432 | 216 | 1288 | 110 | 21 | 99,76 | Eukaryota | Alveolata | Apicomplexa | Apicomplexa_X | Colpodellidae_XX | Colpodellidae_XX+sp. |
| OTU_11 | 0 | 0 | 0 | 0 | 0 | 4 | 3 | 7 | 0 | 0 | 3159 | 95,26 | Eukaryota | Stramenopiles | Ochrophyta | Chrysophyceae-Synurophyceae | Chrysosphaerella | Chrysosphaerella+sp. |
| OTU_12 | 5101 | 4 | 0 | 375 | 64 | 0 | 1 | 52 | 0 | 12 | 104 | 98,33 | Eukaryota | Archaeplastida | Chlorophyta | Trebouxiophyceae | Catena | Catena+viridis |
| OTU_13 | 403 | 0 | 0 | 0 | 0 | 0 | 2238 | 0 | 0 | 392 | 290 | 99,27 | Eukaryota | Cryptista | Cryptophyta | Cryptophyceae | Cryptomonas | Cryptomonas+sp. |
| OTU_14 | 1219 | 243 | 0 | 0 | 1 | 0 | 0 | 119 | 0 | 1 | 41 | 99,31 | Eukaryota | Stramenopiles | Ochrophyta | Eustigmatophyceae | Nannochloropsis | Nannochloropsis+oceanica |
| OTU_15 | 0 | 0 | 0 | 0 | 0 | 0 | 0 | 5 | 1346 | 0 | 0 | 99,06 | Eukaryota | Rhizaria | Filosa | Sarcomonadea | Sandona | Sandona+sp. |
| OTU_16 | 0 | 0 | 0 | 0 | 0 | 0 | 67 | 859 | 0 | 143 | 30 | 100,00 | Eukaryota | Stramenopiles | Ochrophyta | Bacillariophyta | Polar-centric-Mediophyceae_X | Polar-centric-Mediophyceae_X+sp. |
| OTU_17 | 0 | 0 | 8863 | 0 | 3 | 4 | 0 | 0 | 43 | 0 | 2 | 90,74 | Eukaryota | Opisthokonta | Fungi | Chytridiomycota | Rhyzophidiales_X | Rhyzophidiales_X+sp. |
| OTU_18 | 256 | 737 | 1 | 64 | 1674 | 0 | 0 | 5 | 0 | 3 | 7 | 99,52 | Eukaryota | Archaeplastida | Chlorophyta | Trebouxiophyceae | Chloroparva | Chloroparva+pannonica |
| OTU_19 | 0 | 0 | 0 | 0 | 0 | 0 | 765 | 237 | 0 | 32 | 234 | 99,28 | Eukaryota | Archaeplastida | Chlorophyta | Chlorophyceae | Pascherina | Pascherina+tetras |
| OTU_20 | 0 | 0 | 0 | 0 | 0 | 0 | 8 | 790 | 286 | 78 | 106 | 99,03 | Eukaryota | Cryptista | Cryptophyta | Cryptophyceae | Komma | Komma+caudata |
| OTU_22 | 0 | 0 | 0 | 0 | 0 | 2 | 266 | 511 | 0 | 29 | 636 | 98,80 | Eukaryota | Archaeplastida | Chlorophyta | Chlorophyceae | Vitreochlamys | Vitreochlamys+nekrassovii |
| OTU_24 | 1 | 1 | 6571 | 0 | 0 | 1 | 0 | 0 | 0 | 0 | 0 | 99,05 | Eukaryota | Stramenopiles | Ochrophyta | Bacillariophyta | Aulacoseira | Aulacoseira+granulata |
| OTU_25 | 0 | 0 | 0 | 0 | 0 | 1 | 434 | 161 | 0 | 162 | 390 | 98,08 | Eukaryota | Archaeplastida | Chlorophyta | Chlorophyceae | Characiochloris | Characiochloris+sasae |
| OTU_27 | 0 | 0 | 0 | 0 | 2242 | 1 | 0 | 121 | 0 | 71 | 121 | 99,28 | Eukaryota | Cryptista | Katablepharidophyta | Katablepharidaceae | Katablepharidales_XX | Katablepharidales_XX+sp. |
| OTU_28 | 0 | 0 | 0 | 0 | 2222 | 0 | 0 | 0 | 0 | 1 | 7 | 85,05 | Eukaryota | Opisthokonta | Fungi | Basidiomycota | Tremellomycetes_X | Tremellomycetes_X+sp. |
| OTU_29 | 0 | 0 | 3440 | 0 | 1 | 2 | 0 | 6 | 72 | 1 | 2 | 97,84 | Eukaryota | Opisthokonta | Mesomycetozoa | Ichthyosporea | Abeoformidae_Group_MAIP_2_X | Abeoformidae_Group_MAIP_2_X+sp. |
| OTU_31 | 0 | 0 | 0 | 0 | 0 | 0 | 0 | 332 | 14 | 20 | 1 | 97,22 | Eukaryota | Stramenopiles | Stramenopiles_X | MAST | MAST-12_XX | MAST-12_XX+sp. |
| OTU_32 | 0 | 0 | 0 | 0 | 0 | 0 | 0 | 433 | 0 | 5 | 7 | 97,34 | Eukaryota | Stramenopiles | Stramenopiles_X | Bicoecea | Pseudodendromonadales_XX | Pseudodendromonadales_XX+sp. |
| OTU_33 | 0 | 597 | 0 | 0 | 0 | 0 | 0 | 4 | 0 | 0 | 0 | 98,35 | Eukaryota | Stramenopiles | Ochrophyta | Eustigmatophyceae | Monodus | Monodus+subterranea |
| OTU_34 | 0 | 0 | 0 | 0 | 0 | 0 | 0 | 0 | 702 | 0 | 0 | 95,89 | Eukaryota | Alveolata | Apicomplexa | Apicomplexa_X | Colpodellidae_XX | Colpodellidae_XX+sp. |
| OTU_35 | 126 | 0 | 0 | 0 | 0 | 0 | 619 | 4 | 0 | 3 | 0 | 100,00 | Eukaryota | Stramenopiles | Ochrophyta | Bacillariophyta | Chaetoceros | Chaetoceros+sp. |
| OTU_37 | 0 | 0 | 1 | 0 | 0 | 1 | 1 | 20 | 0 | 1 | 1416 | 90,57 | Eukaryota | Stramenopiles | Stramenopiles_X | Bicoecea | Pseudodendromonadales_XX | Pseudodendromonadales_XX+sp. |
| OTU_38 | 0 | 0 | 0 | 0 | 0 | 0 | 1001 | 15 | 50 | 1 | 1 | 97,34 | Eukaryota | Alveolata | Perkinsea | Perkinsida | Perkinsida_XXX | Perkinsida_XXX+sp. |
| OTU_40 | 27 | 0 | 0 | 36 | 0 | 0 | 288 | 13 | 0 | 187 | 265 | 95,63 | Eukaryota | Alveolata | Ciliophora | Spirotrichea | Tintinnidium | Tintinnidium+sp. |
| OTU_41 | 0 | 0 | 0 | 0 | 0 | 0 | 1 | 0 | 0 | 22 | 1462 | 97,71 | Eukaryota | Haptista | Centroheliozoa | Centroheliozoa_X | Pterocystis | Pterocystis+sp. |
| OTU_42 | 0 | 0 | 0 | 0 | 0 | 0 | 1 | 514 | 44 | 14 | 0 | 100,00 | Eukaryota | Stramenopiles | Ochrophyta | Chrysophyceae-Synurophyceae | Pedospumella | Pedospumella+encystans |
| OTU_44 | 0 | 0 | 0 | 0 | 0 | 2 | 0 | 0 | 0 | 2 | 1873 | 84,20 | Eukaryota | Alveolata | Dinophyta | Dinophyceae | Suessiales_XX | Suessiales_XX+sp. |
| OTU_45 | 0 | 0 | 0 | 0 | 0 | 1 | 0 | 16 | 15 | 0 | 1588 | 99,30 | Eukaryota | Haptista | Centroheliozoa | Centroheliozoa_X | Centroheliozoa_XXXX | Centroheliozoa_XXXX+sp. |
| OTU_46 | 0 | 0 | 0 | 0 | 217 | 1 | 5 | 572 | 26 | 186 | 130 | 99,28 | Eukaryota | Archaeplastida | Chlorophyta | Chlorophyceae | Wislouchiella | Wislouchiella+planctonica |
| OTU_47 | 0 | 263 | 0 | 13 | 0 | 0 | 0 | 165 | 13 | 4 | 1 | 99,04 | Eukaryota | Archaeplastida | Chlorophyta | Mamiellophyceae | Monomastix | Monomastix+sp. |
| OTU_48 | 0 | 0 | 0 | 0 | 0 | 0 | 0 | 0 | 568 | 1 | 0 | 88,76 | Eukaryota | Opisthokonta | Fungi | Mucoromycota | Endogone | Endogone+lactiflua |
| OTU_49 | 0 | 0 | 0 | 885 | 0 | 0 | 0 | 0 | 0 | 0 | 0 | 97,85 | Eukaryota | Archaeplastida | Chlorophyta | Trebouxiophyceae | Chlorella | Chlorella+minutissima |
| OTU_50 | 0 | 0 | 0 | 538 | 0 | 0 | 30 | 0 | 0 | 300 | 28 | 95,17 | Eukaryota | Alveolata | Ciliophora | Spirotrichea | Tintinnidium | Tintinnidium+mucicola |
| OTU_51 | 36 | 0 | 0 | 0 | 0 | 0 | 98 | 396 | 5 | 23 | 11 | 97,37 | Eukaryota | Alveolata | Apicomplexa | Apicomplexa_X | Colpodellidae_XX | Colpodellidae_XX+sp. |
| OTU_52 | 0 | 0 | 0 | 0 | 0 | 0 | 2 | 416 | 0 | 7 | 0 | 96,20 | Eukaryota | Alveolata | Perkinsea | Perkinsida | Perkinsida_XXX | Perkinsida_XXX+sp. |
| OTU_55 | 0 | 0 | 0 | 0 | 0 | 0 | 82 | 237 | 0 | 230 | 3 | 100,00 | Eukaryota | Stramenopiles | Ochrophyta | Bacillariophyta | Thalassiosira | Thalassiosira+pseudonana |
| OTU_56 | 0 | 0 | 0 | 0 | 0 | 0 | 532 | 37 | 0 | 362 | 275 | 98,56 | Eukaryota | Archaeplastida | Chlorophyta | Chlorophyceae | Chlamydomonas | Chlamydomonas+sp. |
| OTU_57 | 0 | 0 | 0 | 0 | 0 | 0 | 0 | 357 | 5 | 12 | 0 | 97,37 | Eukaryota | Alveolata | Dinophyta | Dinophyceae | Prorocentrum | Prorocentrum+sp. |
| OTU_58 | 202 | 86 | 138 | 29 | 1337 | 0 | 0 | 0 | 0 | 3 | 0 | 96,89 | Eukaryota | Archaeplastida | Chlorophyta | Trebouxiophyceae | Chlorella | Chlorella+vulgaris |
| OTU_59 | 1300 | 0 | 0 | 0 | 0 | 0 | 30 | 37 | 0 | 13 | 0 | 94,00 | Eukaryota | Opisthokonta | Fungi | Fungi_X | Fungi_XXXX | Fungi_XXXX+sp. |
| OTU_60 | 0 | 0 | 0 | 0 | 0 | 2 | 9 | 98 | 0 | 0 | 513 | 87,03 | Eukaryota | Opisthokonta | Fungi | Fungi_X | Fungi_XXXX | Fungi_XXXX+sp. |
| OTU_61 | 0 | 0 | 0 | 0 | 0 | 0 | 719 | 0 | 0 | 1 | 0 | 99,52 | Eukaryota | Archaeplastida | Chlorophyta | Chlorophyceae | CW-Chlamydomonadales_X | CW-Chlamydomonadales_X+sp. |
| OTU_63 | 719 | 0 | 0 | 385 | 8 | 0 | 11 | 16 | 0 | 6 | 1 | 99,28 | Eukaryota | Archaeplastida | Chlorophyta | Chlorophyceae | Rhombocystis | Rhombocystis+complanata |
| OTU_64 | 57 | 477 | 0 | 0 | 187 | 0 | 0 | 1 | 0 | 0 | 0 | 97,13 | Eukaryota | Archaeplastida | Chlorophyta | Trebouxiophyceae | Meyerella | Meyerella+planktonica |
| OTU_65 | 74 | 0 | 0 | 0 | 0 | 0 | 7 | 304 | 13 | 5 | 13 | 99,54 | Eukaryota | Stramenopiles | Stramenopiles_X | Oomycota | Peronosporales_X | Peronosporales_X+sp. |
| OTU_67 | 0 | 0 | 1 | 0 | 0 | 0 | 0 | 384 | 0 | 1 | 15 | 94,74 | Eukaryota | Archaeplastida | Chlorophyta | Trebouxiophyceae | Micractinium | Micractinium+pusillum |
| OTU_70 | 0 | 114 | 1979 | 0 | 0 | 0 | 0 | 46 | 21 | 0 | 1 | 98,00 | Eukaryota | Opisthokonta | Fungi | Chytridiomycota | Chytridiomycetes_X | Chytridiomycetes_X+sp._strain9 |
| OTU_72 | 0 | 0 | 0 | 0 | 0 | 0 | 534 | 0 | 0 | 73 | 16 | 99,52 | Eukaryota | Archaeplastida | Chlorophyta | Chlorophyceae | Chlamydomonas | Chlamydomonas+sp. |
| OTU_73 | 0 | 0 | 0 | 0 | 0 | 0 | 0 | 0 | 1491 | 0 | 0 | 94,12 | Eukaryota | Opisthokonta | Choanoflagellida | Choanoflagellatea | Codosiga | Codosiga+sp. |
| OTU_74 | 0 | 0 | 0 | 0 | 0 | 1 | 363 | 48 | 0 | 5 | 1 | 99,28 | Eukaryota | Stramenopiles | Ochrophyta | Chrysophyceae-Synurophyceae | Clade-C_X | Clade-C_X+sp. |
| OTU_75 | 0 | 273 | 0 | 0 | 0 | 0 | 7 | 230 | 0 | 1 | 14 | 99,04 | Eukaryota | Archaeplastida | Chlorophyta | Chlorophyceae | Sphaeropleales_X | Sphaeropleales_X+sp. |
| OTU_76 | 26 | 22 | 0 | 5 | 0 | 0 | 174 | 59 | 0 | 312 | 105 | 98,81 | Eukaryota | Stramenopiles | Ochrophyta | Bacillariophyta | Skeletonema | Skeletonema+grethae |
| OTU_77 | 0 | 0 | 0 | 0 | 0 | 0 | 137 | 0 | 0 | 483 | 3 | 98,56 | Eukaryota | Rhizaria | Filosa | Imbricatea | Novel-clade-2_X | Novel-clade-2_X+sp. |
| OTU_81 | 0 | 0 | 0 | 0 | 49 | 0 | 279 | 11 | 16 | 0 | 8 | 99,77 | Eukaryota | Rhizaria | Filosa | Thecofilosea | Mataza-lineage_X | Mataza-lineage_X+sp. |
| OTU_82 | 0 | 0 | 0 | 0 | 368 | 0 | 0 | 95 | 0 | 21 | 50 | 97,85 | Eukaryota | Stramenopiles | Ochrophyta | Bacillariophyta | Raphid-pennate_X | Raphid-pennate_X+sp. |
| OTU_84 | 0 | 0 | 0 | 0 | 1 | 0 | 184 | 80 | 0 | 231 | 259 | 100,00 | Eukaryota | Archaeplastida | Chlorophyta | Chlorophyceae | Chlamydomonas | Chlamydomonas+sp. |
| OTU_87 | 1698 | 182 | 0 | 142 | 96 | 1 | 0 | 1 | 0 | 0 | 0 | 98,09 | Eukaryota | Archaeplastida | Chlorophyta | Trebouxiophyceae | Chlorella | Chlorella+sp. |
| OTU_88 | 0 | 0 | 452 | 0 | 0 | 0 | 0 | 19 | 135 | 24 | 3 | 95,48 | Eukaryota | Rhizaria | Cercozoa | Filosa-Imbricatea | Novel-clade-2_X | Novel-clade-2_X+sp._strain15 |
| OTU_89 | 607 | 16 | 0 | 0 | 0 | 152 | 54 | 8 | 3 | 4 | 27 | 99,52 | Eukaryota | Alveolata | Ciliophora | Spirotrichea | Halteria | Halteria+grandinella |
| OTU_91 | 0 | 0 | 0 | 0 | 294 | 0 | 0 | 58 | 0 | 0 | 123 | 99,76 | Eukaryota | Stramenopiles | Ochrophyta | Bacillariophyta | Synedra | Synedra+berolinensis |
| OTU_92 | 0 | 0 | 0 | 0 | 65 | 1 | 0 | 61 | 0 | 0 | 272 | 92,74 | Eukaryota | Stramenopiles | Ochrophyta | Chrysophyceae-Synurophyceae | Chrysophyceae-Synurophyceae_XXX | Chrysophyceae-Synurophyceae_XXX+sp. |
| OTU_102 | 248 | 0 | 0 | 0 | 0 | 0 | 74 | 1 | 0 | 0 | 0 | 91,98 | Eukaryota | Stramenopiles | Ochrophyta | Bacillariophyta | Hemiaulus | Hemiaulus+sinensis |
| OTU_103 | 73 | 345 | 0 | 0 | 0 | 0 | 0 | 0 | 0 | 0 | 0 | 94,50 | Eukaryota | Archaeplastida | Chlorophyta | Pedinophyceae | Chlorochytridion | Chlorochytridion+sp. |
| OTU_105 | 0 | 0 | 0 | 0 | 0 | 0 | 0 | 8 | 243 | 0 | 9 | 100,00 | Eukaryota | Stramenopiles | Ochrophyta | Chrysophyceae-Synurophyceae | Synurales_XX | Synurales_XX+sp. |
| OTU_111 | 0 | 0 | 0 | 0 | 269 | 0 | 22 | 71 | 0 | 22 | 84 | 99,04 | Eukaryota | Stramenopiles | Ochrophyta | Bacillariophyta | Amphora | Amphora+sp. |
| OTU_114 | 0 | 0 | 0 | 298 | 302 | 0 | 0 | 0 | 0 | 10 | 0 | 96,65 | Eukaryota | Archaeplastida | Chlorophyta | Trebouxiophyceae | Nannochloris | Nannochloris+sp. |
| OTU_115 | 423 | 1 | 0 | 0 | 0 | 0 | 0 | 0 | 0 | 0 | 0 | 96,66 | Eukaryota | Opisthokonta | Fungi | Chytridiomycota | Spizellomyces | Spizellomyces+palustris |
| OTU_116 | 0 | 427 | 0 | 0 | 1 | 0 | 33 | 1 | 0 | 1 | 0 | 93,75 | Eukaryota | Archaeplastida | Chlorophyta | Pedinophyceae | Chlorochytridion | Chlorochytridion+tuberculatum |
| OTU_121 | 0 | 0 | 0 | 0 | 0 | 0 | 0 | 0 | 0 | 0 | 420 | 88,94 | Eukaryota | Opisthokonta | Choanoflagellida | Choanoflagellida_X | Sphaeroeca | Sphaeroeca+volvox |
| OTU_126 | 0 | 0 | 59 | 0 | 0 | 0 | 0 | 0 | 356 | 0 | 0 | 99,76 | Eukaryota | Opisthokonta | Fungi | Blastocladiomycota | Blastocladiomycota_XX | Blastocladiomycota_XX+sp. |
| OTU_128 | 0 | 0 | 0 | 20 | 395 | 1 | 0 | 85 | 0 | 329 | 412 | 99,29 | Eukaryota | Stramenopiles | Ochrophyta | Bacillariophyta | Cyclostephanos | Cyclostephanos+dubius |
| OTU_129 | 20 | 0 | 0 | 2304 | 11 | 0 | 0 | 0 | 0 | 14 | 0 | 98,56 | Eukaryota | Archaeplastida | Chlorophyta | Trebouxiophyceae | Chlorellales_XX | Chlorellales_XX+sp._strain4 |
| OTU_130 | 1484 | 0 | 0 | 0 | 3 | 0 | 718 | 215 | 63 | 234 | 128 | 99,05 | Eukaryota | Stramenopiles | Ochrophyta | Bacillariophyta | Cyclotella | Cyclotella+meneghiniana |
| OTU_135 | 0 | 0 | 0 | 0 | 0 | 0 | 110 | 0 | 23 | 542 | 109 | 98,42 | Eukaryota | Haptista | Centroheliozoa | Centroheliozoa_X | Heterophrys | Heterophrys+sp. |
| OTU_144 | 0 | 0 | 0 | 0 | 0 | 2 | 6 | 0 | 0 | 0 | 303 | 87,15 | Eukaryota | Rhizaria | Endomyxa | Proteromyxida | Vampyrella | Vampyrella+sp. |
| OTU_152 | 0 | 0 | 0 | 0 | 0 | 393 | 0 | 0 | 0 | 0 | 0 | 91,16 | Eukaryota | Rhizaria | Cercozoa | Filosa | Filosa_XXX | Filosa_XXX+sp. |
| OTU_153 | 0 | 0 | 0 | 0 | 0 | 0 | 0 | 0 | 0 | 598 | 0 | 99,76 | Eukaryota | Stramenopiles | Ochrophyta | Chrysophyceae-Synurophyceae | Chrysophyceae-Synurophyceae_XXX | Chrysophyceae-Synurophyceae_XXX+sp. |
| OTU_163 | 0 | 0 | 0 | 0 | 0 | 0 | 0 | 6 | 0 | 517 | 11 | 97,85 | Eukaryota | Stramenopiles | Ochrophyta | Bacillariophyta | Bacillariophyta | Bacillariophyta+sp. |
| OTU_170 | 0 | 0 | 0 | 0 | 0 | 0 | 0 | 0 | 366 | 0 | 0 | 94,12 | Eukaryota | Opisthokonta | Choanoflagellida | Choanoflagellatea | Codosiga | Codosiga+sp. |
| OTU_171 | 0 | 59 | 0 | 0 | 268 | 0 | 0 | 0 | 0 | 0 | 0 | 98,09 | Eukaryota | Archaeplastida | Chlorophyta | Trebouxiophyceae | Chlorella | Chlorella+vulgaris |
| OTU_176 | 0 | 0 | 0 | 0 | 0 | 0 | 20 | 20 | 0 | 62 | 282 | 97,60 | Eukaryota | Archaeplastida | Chlorophyta | Chlorophyceae | CW-Chlamydomonadales_X | CW-Chlamydomonadales_X+sp. |
| OTU_179 | 0 | 0 | 448 | 0 | 0 | 0 | 0 | 0 | 0 | 0 | 1 | 91,38 | Eukaryota | Rhizaria | Filosa | Sarcomonadea | Eocercomonas | Eocercomonas+sp. |
| OTU_191 | 0 | 0 | 0 | 0 | 5 | 11 | 0 | 100 | 6 | 249 | 76 | 98,95 | Eukaryota | Opisthokonta | Fungi | Chytridiomycota | Chytridiomycetes_X | Chytridiomycetes_X+sp._strain3 |
| OTU_192 | 0 | 0 | 0 | 0 | 0 | 0 | 0 | 0 | 0 | 623 | 128 | 99,52 | Eukaryota | Archaeplastida | Chlorophyta | Chlorophyceae | CW-Chlamydomonadales_X | CW-Chlamydomonadales_X+sp. |
| OTU_202 | 0 | 0 | 0 | 0 | 0 | 0 | 0 | 0 | 0 | 289 | 0 | 99,00 | Eukaryota | Alveolata | Ciliophora | Heterotrichea | Climacostomum | Climacostomum+virens |
| OTU_207 | 0 | 0 | 437 | 0 | 0 | 0 | 0 | 0 | 0 | 0 | 0 | 94,47 | Eukaryota | Archaeplastida | Chlorophyta | Trebouxiophyceae | Oocystaceae | Oocystaceae+sp. |
| OTU_252 | 0 | 0 | 0 | 0 | 0 | 0 | 0 | 0 | 470 | 0 | 0 | 94,12 | Eukaryota | Opisthokonta | Choanoflagellida | Choanoflagellatea | Codosiga | Codosiga+sp. |
| OTU_333 | 0 | 0 | 0 | 0 | 0 | 0 | 0 | 0 | 0 | 273 | 0 | 95,88 | Eukaryota | Alveolata | Perkinsea | Perkinsida | Perkinsida_XXX | Perkinsida_XXX+sp. |
| OTU_356 | 0 | 0 | 0 | 0 | 0 | 0 | 0 | 415 | 0 | 5 | 2 | 96,37 | Eukaryota | Alveolata | Perkinsea | Perkinsida | Perkinsida_XXX | Perkinsida_XXX+sp. |
| OTU_391 | 0 | 0 | 0 | 0 | 0 | 0 | 0 | 0 | 0 | 4 | 257 | 98,80 | Eukaryota | Archaeplastida | Chlorophyta | Chlorophyceae | Chlamydomonas | Chlamydomonas+subcaudata |
| OTU_526 | 1816 | 78 | 3 | 120 | 792 | 0 | 7 | 1 | 0 | 3 | 1 | 97,61 | Eukaryota | Archaeplastida | Chlorophyta | Trebouxiophyceae | Catena | Catena+viridis |
| OTU_572 | 0 | 0 | 0 | 0 | 0 | 0 | 0 | 0 | 0 | 463 | 5 | 98,08 | Eukaryota | Archaeplastida | Chlorophyta | Chlorophyceae | Chlamydomonas | Chlamydomonas+sordida |
| OTU_576 | 0 | 0 | 0 | 0 | 0 | 0 | 0 | 83 | 0 | 75 | 264 | 96,88 | Eukaryota | Archaeplastida | Chlorophyta | Chlorophyceae | Chlamydomonas | Chlamydomonas+sp. |
| OTU_582 | 0 | 0 | 0 | 0 | 670 | 0 | 0 | 199 | 0 | 0 | 25 | 98,56 | Eukaryota | Archaeplastida | Chlorophyta | Trebouxiophyceae | Nannochloris | Nannochloris+sp. |

La Juanita (LJ); Chascomús (CH); La Limpia (LI); Mar Chiquita (MA2); Vitel (VI); Los Quilmes (QU);

Supplementary Table 8: Bray-Curtis disimilarity index for sorting samples.

| | LJ | CH | LI | MA2 | VI |
|---|---|---|---|---|---|
| CH | 0.75 | - | - | - | - |
| LI | 0.91 | 0.89 | - | - | - |
| MA2 | 0.73 | 0.63 | 0.91 | - | - |
| VI | 0.75 | 0.64 | 0.91 | 0.63 | - |
| QU | 0.87 | 0.78 | 0.90 | 0.85 | 0.82 |

La Juanita (LJ); Chascomús (CH); La Limpia (LI); Mar Chiquita (MA2); Vitel (VI); Los Quilmes (QU).
